# An optimised system for rapid auxin-inducible protein degradation in budding yeast

**DOI:** 10.1101/2025.09.24.678194

**Authors:** Petra Hubbe, Charu Sharma, Oliver Pajonk, Niklas Peters, Nadja Guschtschin-Schmidt, Natalie Friemel, Sebastian Schuck

**Affiliations:** Heidelberg University Biochemistry Center, 69120 Heidelberg, Germany; Immundiagnostik AG, 64625 Bensheim, Germany

## Abstract

The auxin system for inducible protein degradation is a powerful tool to investigate protein function. It consists of a degron fused to a target protein, an auxin-related ligand that binds to the degron, and a receptor that recognises the auxin-bound degron and mediates proteasomal degradation of the target protein. Variants of all system components are available, and we here test three degrons, three auxins and three degron receptors to identify optimal combinations of these variants in budding yeast. We show that the degrons mIAA7 or AID* together with adamantyl-auxin and the degron receptor OsTIR1(F74G) allow particularly rapid and extensive degradation. Basal degradation in the absence of auxin is generally low and can be prevented entirely by inducible expression of OsTIR1(F74G). Finally, we demonstrate that the remarkable efficiency of this system makes it competitive with established chemical inhibitors, such as tunicamycin and MG132, and with temperature-sensitive mutant alleles. These findings will aid the effective application of the auxin system.

## Introduction

Biological research often aims to define the physiological functions of individual proteins. The most direct approach for this purpose is to inactivate a protein of interest and examine the resulting cellular or organismal phenotypes. Ideally, protein inactivation should be specific, complete and immediate. Non-essential proteins can be removed through knockout of the corresponding genes, which is specific and complete, but slow. It takes many cell division cycles until newly created mutant cells can be analysed so that cell adaptation may gradually obscure the knockout phenotype. Similar limitations apply when the mRNA of a target protein is eliminated, for instance through RNA interference, because the impact on protein function depends on how quickly pre-existing protein molecules are degraded. If a target protein is long-lived, its levels will decline only slowly. In the meantime, cell adaptation may again alter the observed phenotype. Given these caveats, target protein inactivation should be as fast as possible.

Several systems are available for rapid target protein inactivation in eukaryotic cells by inducible mislocalisation or degradation (Prozzillo et al, 2020; Bondeson et al, 2022; Hatoyama et al, 2024). The dTag and BromoTag systems require the introduction of two exogenous components into cells: a degradation determinant (called degron), which is fused to the target protein by genome editing, and a chemical inducer, which binds to the degron and mediates degron recognition by the ubiquitin-proteasome system (Nabet et al, 2018; Bond et al, 2021). The auxin system requires a third exogenous component, namely a protein that bridges the inducer-bound degron and the ubiquitin-proteasome system (Nishimura et al, 2009). Currently, the auxin system is the only option for studies in budding yeast, *Saccharomyces cerevisiae*, because no degron is known that is directly recognised by the yeast ubiquitin-proteasome system in an inducer-dependent manner.

The auxin system is based on the plant hormone indole-3-acetic acid, which binds to Aux/IAA proteins and mediates their interaction with F-box proteins of the TIR1/AFB family. In this way, indole-3-acetic acid triggers recognition of Aux/IAA proteins by TIR1/AFB-containing ubiquitin ligase complexes and subsequent degradation of Aux/IAA by the proteasome (Leyser, 2018). The system can be transferred to non-plant cells. For this, a TIR1/AFB protein is introduced into the cells of interest and an Aux/IAA protein is fused to a target protein. Aux/IAA then serves as a conditional degron so that the target protein is degraded upon addition of indole-3-acetic acid (Nishimura et al, 2009). This transfer across species is possible because TIR1/AFB proteins are incorporated into SCF ubiquitin ligase complexes in many different organisms (Phanindhar and Mishra, 2023).

All components of the auxin system have been improved through engineering efforts, yielding different Aux/IAA variants (degrons), indole-3-acetic acid derivatives (auxins) and TIR1/AFB variants (degron receptors). These improvements have reduced degron size, restricted basal degradation in the absence of auxin, lowered the auxin concentration and thus avoided off-target effects, and increased the speed of target protein degradation. A key advance was the implementation of a bump-and-hole strategy, in which enlargement of auxin through addition of bulky chemical moieties and corresponding enlargement of the auxin binding pocket of degron receptors through amino acid exchanges strengthened the interaction between auxin-bound degron and receptor (Uchida et al, 2018; Yamada et al, 2018; Nishimura et al, 2020). Various system components are now available, raising the question which ones should be combined for optimal target protein degradation.

Here, we systematically evaluated combinations of three degrons, three auxins and three degron receptors in budding yeast (Figure 1A). The degrons were the shortened AtIAA17 variants mini-AID (mAID; Kubota et al, 2013) and AID* (Murowska and Ulrich, 2013), and the shortened AtIAA7 variant mini-IAA7 (mIAA7; Li et al, 2019). The inducers were the natural indole-3-acetic acid and its artificial derivatives 5-phenyl- and 5-adamantyl-indole- 3-acetic acid (Uchida et al, 2018; Yamada et al, 2018). We call these variants n-auxin, p- auxin and a-auxin, respectively. The degron receptors were the natural OsTIR1 and the OsTIR1(F74G) and AtAFB(F74A) variants with enlarged auxin binding pockets (Nishimura et al, 2009; Yesbolatova et al, 2020; Li et al, 2024). We show that the combination of mIAA7, a-auxin and OsTIR1(F74G) allows extensive and rapid inducible protein degradation, with minimal basal degradation. Furthermore, we demonstrate that this optimised system is competitive with established chemical inhibitors and temperature-sensitive mutant alleles.

**Figure 1.**
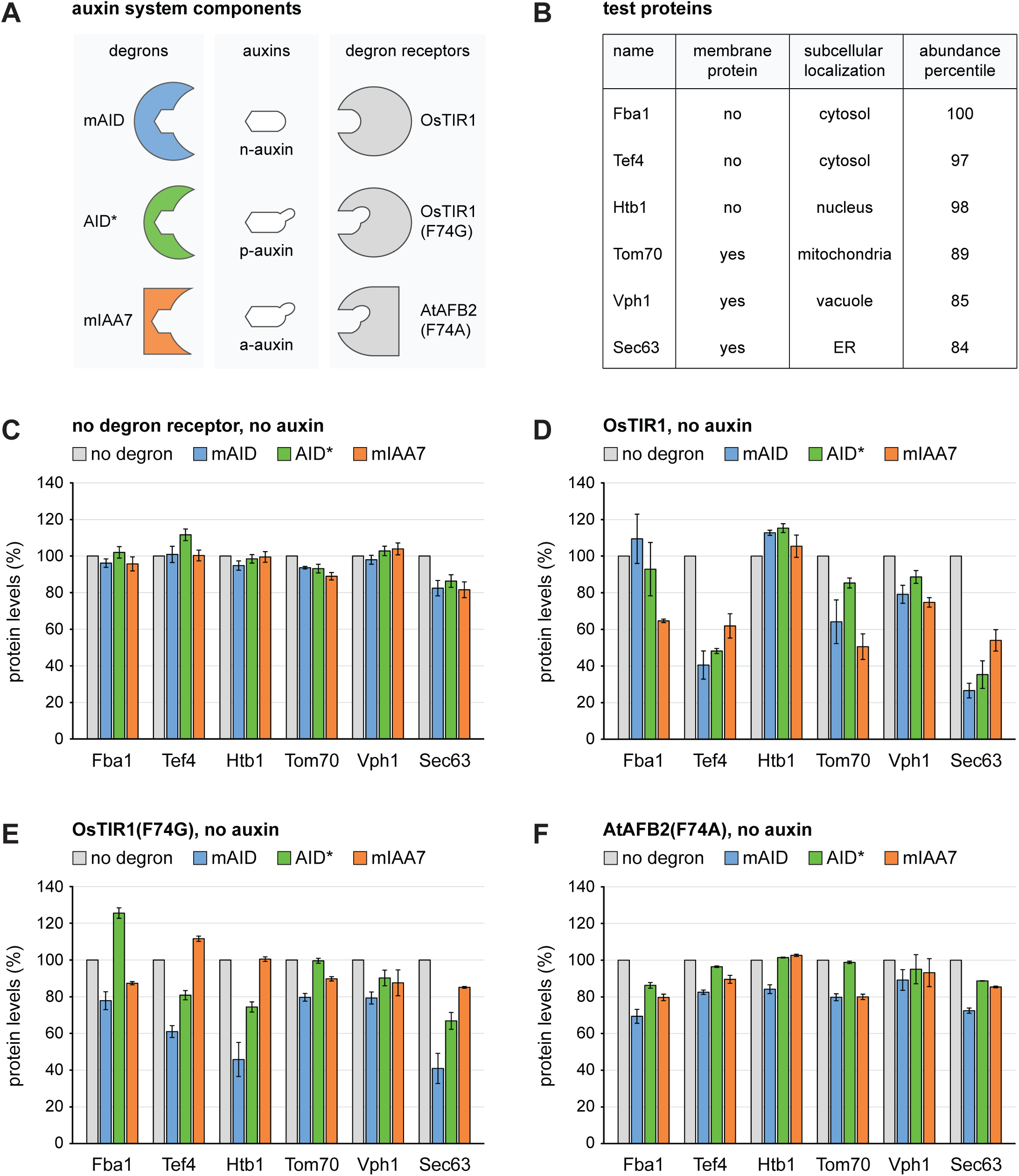
Auxin system components, test proteins and basal degradation in the absence of auxin. **(A)** Degrons, auxins and degron receptors used in this study. mAID (8 kDa) and AID* (5 kDa) are derived from AtIAA17, mIAA7 (8 kDa) is derived from AtIAA7. N-auxin is the naturally occurring indole-3-acetic acid, p-auxin and a-auxin contain an additional phenyl and adamantyl moiety, respectively. OsTIR1 is a naturally occurring degron receptor, OsTIR1(F74G) and AtAFB2(F74A) are mutant variants in which the auxin binding pocket has been enlarged to accommodate the artificial auxins p-auxin and a- auxin. **(B)** Test proteins used to evaluate combinations of degron, auxin and degron receptor. Cellular levels of these proteins are in the 84^th^ to 100^th^ abundance percentile of all yeast proteins detectable by mass spectrometry (Platzek et al, 2025). **(C)** Flow cytometry measurement of relative levels of test proteins tagged with mNeonGreen (no degron), mAID-mNeonGreen, AID*-mNeonGreen or mIAA7-mNeonGreen. Cells did not express a degron receptor and were not exposed to auxin. Bars show the mean protein levels of at least three biological replicates, error bars show the standard error of the mean. Protein levels were normalised to the levels in cells expressing the respective test protein tagged only with mNeonGreen. **(D-F)** As in panel C but in cells expressing OsTIR1, OsTIR1(F74G) and AtAFB2(F74A), respectively.

## Results

### Basal protein degradation by degron tagging and expression of degron receptors

To assess combinations of degrons, auxins and degron receptors, we determined basal and auxin-inducible degradation of six test proteins. This set consisted of soluble and transmembrane proteins from different subcellular compartments. Specifically, we chose (1) the cytosolic glycolytic enzyme Fba1, (2) the cytosolic translation elongation factor Tef4, (3) the nuclear histone Htb1, (4) the mitochondrial outer membrane protein Tom70, (5) the vacuolar membrane protein Vph1, and (6) the endoplasmic reticulum (ER) membrane protein Sec63 (Figure 1B). The localisations and topologies of these proteins ensure that degrons fused to their C-termini are exposed to the cytosol or nucleoplasm, and are thus accessible to the ubiquitin-proteasome system. Furthermore, all six proteins are among the 20% most abundant proteins in yeast (Platzek et al, 2025). These high levels were chosen to challenge the capacity of the auxin-inducible degradation system.

We first asked whether fusion of the degrons to the test proteins caused degradation in the absence of auxins or degron receptors. We used chromosomal gene tagging to generate strains in which the C-termini of test proteins were fused to the fluorescent protein mNeonGreen or to degron-mNeonGreen cassettes containing mAID, AID* or mIAA7. Quantification of cellular mNeonGreen fluorescence by flow cytometry showed that the test proteins were not destabilised by the degrons. An exception was Sec63, whose levels were lowered by 20% upon fusion to degron-mNeonGreen cassettes compared with fusion to mNeonGreen alone (Figure 1C). In addition, fusion of Htb1 to mAID-mNeonGreen yielded yeast with a severe growth defect, indicating that Htb1-mAID- mNeonGreen was functionally impaired. We then tested whether auxins changed the abundance of degron-tagged test proteins in the absence of a degron receptor, but they did not (Figure S1). Hence, tagging a target protein with one of the degrons is unlikely to cause destabilisation in the absence of a degron receptor, even upon auxin treatment. However, there may be exceptions and the tagged protein may not be fully functional.

Next, we determined whether expression of any of the three degron receptors led to basal degradation of degron-tagged proteins in the absence of auxins. Expression of OsTIR1 under the constitutive *ADH* promoter had only minor destabilising effects on degron-tagged Fba1, Htb1 and Vph1. However, OsTIR1 reduced the levels of degron-tagged Tef4, Tom70 and Sec63 to varying extents, which could exceed 50% (Figure 1D). These results are consistent with previously reported basal degradation of degron-tagged proteins by OsTIR1, which likely arises from auxin-independent association of OsTIR1 with auxin-binding degrons (Yesbolatova et al, 2020). Expression of OsTIR1(F74G) or AtAFB2(F74A) had smaller destabilising effects on degron-tagged test proteins (Figure 1E, F). Hence, these degron receptor variants alleviate the problem of basal degradation, as reported (Yesbolatova et al, 2020; Li et al, 2024). However, the mAID degron together with OsTIR1(F74G) still caused substantial basal degradation of several test proteins (Figure 1E). The AID* and mIAA7 degrons in combination with OsTIR1(F74G) and AtAFB2(F74A) generally caused only minor basal degradation. We therefore excluded OsTIR1 from further analysis and focused on OsTIR1(F74G) and AtAFB2(F74A).

### Auxin-inducible protein degradation by different combinations of system components

We then turned to auxin-induced protein degradation and compared the different auxins in end-point measurements. Guided by previous studies, we used n-auxin at 375 µM, p- auxin at 5 µM and a-auxin at 0.5 µM (Yesbolatova et al, 2020; Yamada et al, 2018). The extent of inducible degradation after two hours differed between test proteins and also depended on the degron and the degron receptor (Figures 2 and S2). For example, Fba1, which belongs to the ten most abundant proteins in yeast (Platzek et al, 2025), showed only moderate degradation. By contrast, Sec63 levels were reduced by at least 80%, regardless of the degron and degron receptor used. However, across all proteins, p-auxin and a-auxin yielded more extensive degradation than n-auxin. In addition, a-auxin performed as well or better than p-auxin, even though its concentration was ten-fold lower. We therefore only used a-auxin in subsequent experiments.

**Figure 2.**
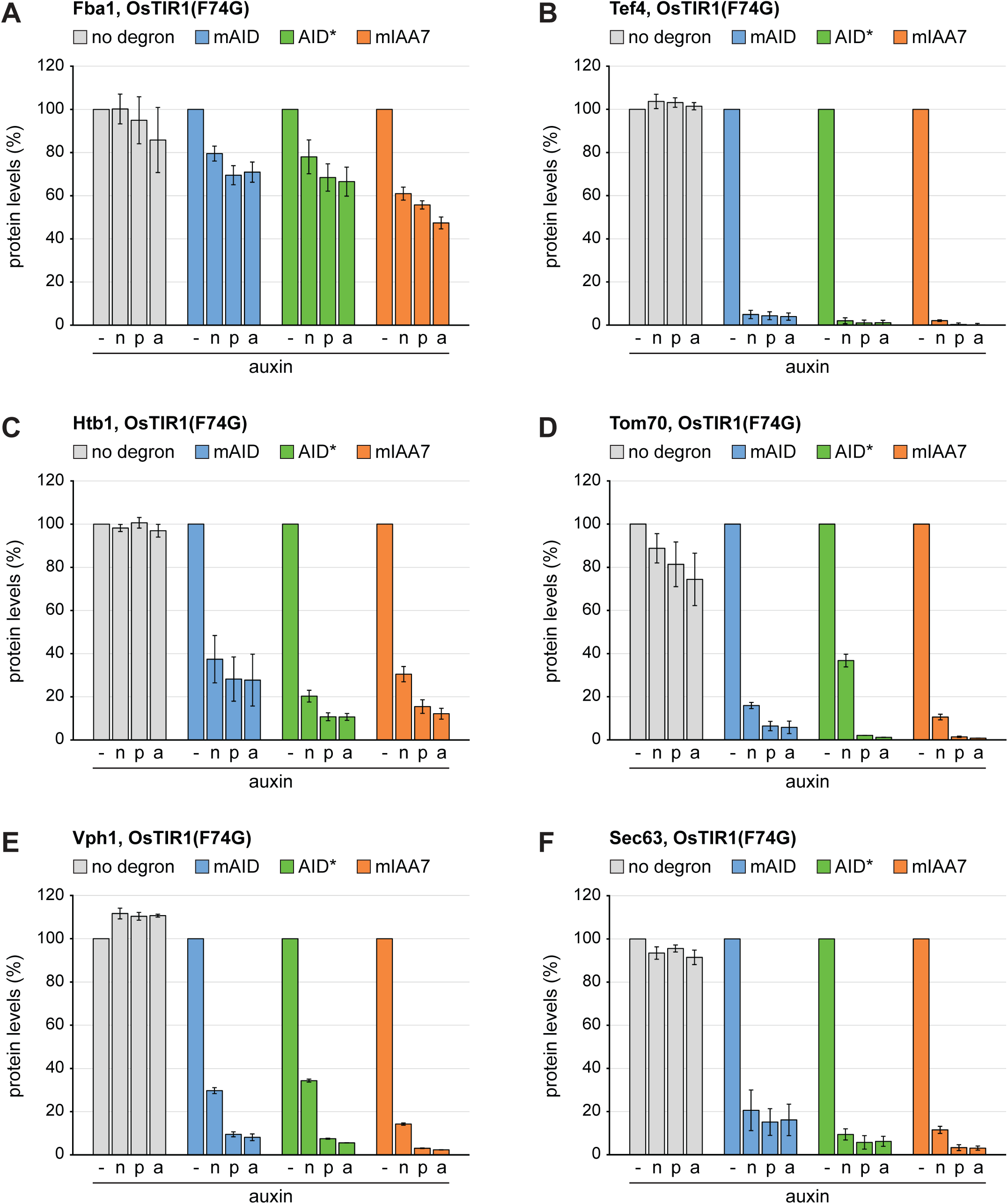
Inducible protein degradation by n-/p-/a-auxin in the presence of OsTIR1(F74G). **(A)** Flow cytometry measurement of relative levels of Fba1 tagged with mNeonGreen (no degron), mAID-mNeonGreen (mAID), AID*-mNeonGreen (AID*) or mIAA7-mNeonGreen (mIAA7). Cells expressed OsTIR1(F74G) and were either not treated or treated with 375 µM n-auxin, 5 µM p-auxin or 0.5 µM a-auxin for 120 minutes. Bars show the mean protein levels of three biological replicates, error bars show the standard error of the mean. Protein levels were normalised to the levels in untreated cells. **(B-F)** As in panel A but for Tef4, Htb1, Tom70, Vph1 and Sec63, respectively.

As a next step, we determined which degron and degron receptor worked best with a- auxin. The end-point measurements above suggested that OsTIR1(F74G) was superior to AtAFB2(F74A). The point mutations OsTIR1(F74A) and AtAFB2(F74G) have also been tested (Yesbolatova et al, 2020; Li et al, 2024) and we confirmed that they offered no advantage over OsTIR1(F74G) and AtAFB2(F74A), respectively (Figure S3). We then carried out time-course measurements to follow degradation of degron-tagged test proteins by a-auxin in the presence of OsTIR1(F74G) or AtAFB2(F74A). For each protein, we normalised its fluorescence in strains with various degrons to its fluorescence in strains in which it was tagged only with mNeonGreen. The resulting plots visualise basal degradation as differences between protein levels at the 0-minute time point and show auxin-induced degradation as decline in protein levels up to the 120-minute time point (Figures 3 and S4). As expected, degradation of Fba1 was inefficient. Depletion was at most 50%, which was achieved by OsTIR1(F74G) together with mAID or mIAA7 (Figure S4A, B). The other proteins were degraded more efficiently and degradation rates correlated inversely with protein abundance. Specifically, the order of degradation rates was Htb1 < Tef4 << Vph1 < Tom70 ≈ Sec63, while the order of abundance was Htb1 > Tef4 >> Tom70 > Vph1 ≈ Sec63. Furthermore, OsTIR1(F74G) yielded faster degradation than AtAFB2(F74A) for every protein and degron. The performance of the three degrons in the presence of OsTIR1(F74G) was similar but mIAA7 showed the least basal degradation (Tef4, Htb1, Sec63 in Figures 3A, S4C and 3E; also see Figure 1E). Thus, mIAA7 together with OsTIR1(F74G) provided the best combination of low basal degradation and rapid auxin-inducible degradation. Remarkably, a-auxin treatment of cells expressing OsTIR1(F74G) reduced the levels of mIAA7-tagged Tom70, Vph1 and Sec63 by approximately 90% within 30 minutes and by >95% within 60 minutes (Figures S4E, 3C and 3E). Even Tef4, which is among the one hundred most abundant proteins in yeast (out of about 5400 proteins present; Ho et al, 2018), was depleted by >95% within 120 minutes (Figure 3A).

**Figure 3.**
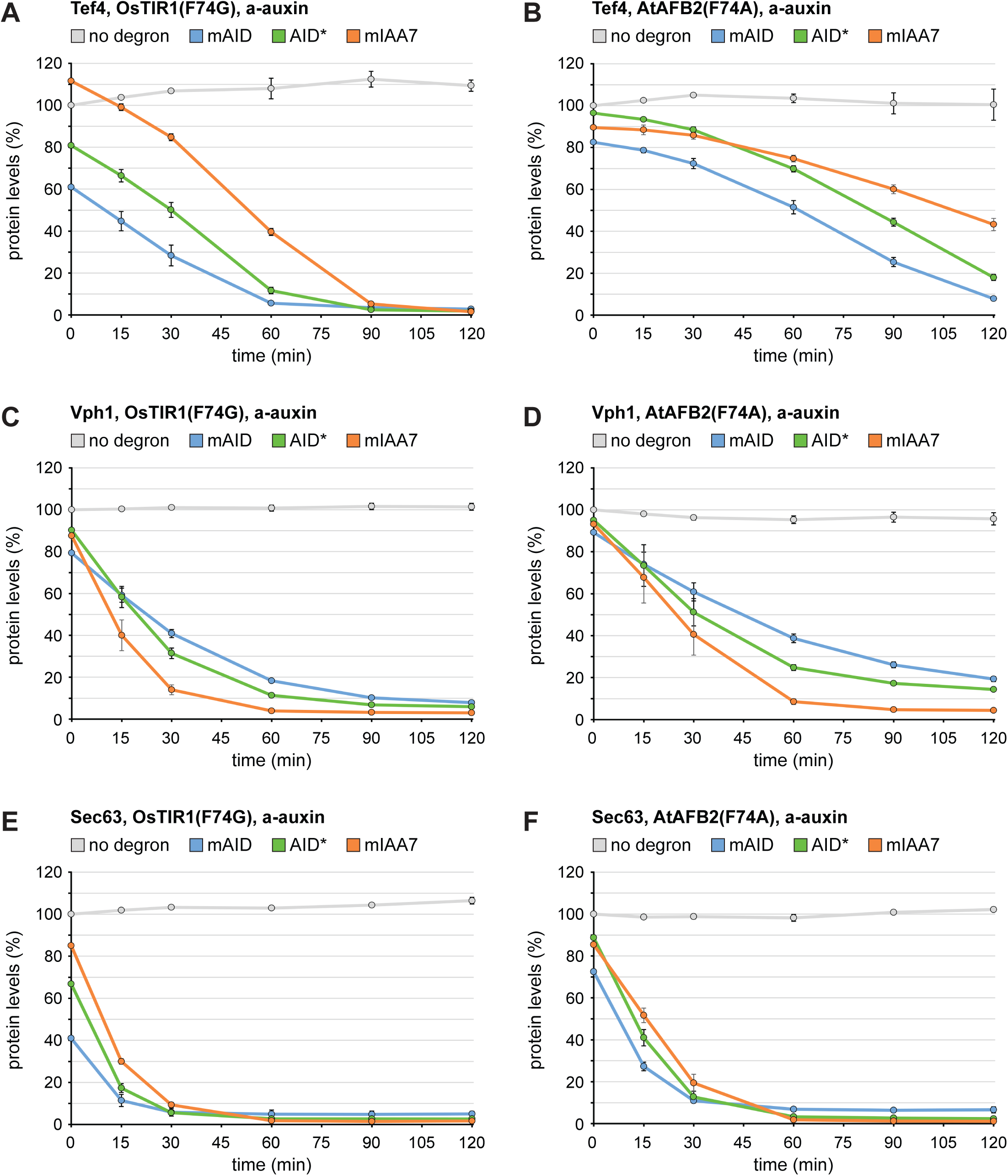
Kinetics of inducible protein degradation by a-auxin in the presence of OsTIR1(F74G) or AtAFB2(F74A). **(A)** Flow cytometry measurement of relative levels of Tef4 tagged with mNeonGreen (no degron), mAID-mNeonGreen (mAID), AID*- mNeonGreen (AID*) or mIAA7-mNeonGreen (mIAA7). Cells expressed OsTIR1(F74G) and were treated with 0.5 µM a-auxin for 0, 15, 30, 60, 90 or 120 minutes. Lines show the mean protein levels of three biological replicates, error bars show the standard error of the mean. Protein levels were normalised to the levels in cells that expressed Tef4- mNeonGreen and were treated with a-auxin for 0 min. **(B)** As in panel A but for cells expressing AtAFB2(F74A). **(C, E)** As in panel A but for cells expressing tagged variants of Vph1 and Sec63, respectively. **(D, F)** As in panel A but for cells expressing AtAFB2(F74A) and tagged variants of Vph1 and Sec63, respectively.

In conclusion, the three degrons together with a-auxin and OsTIR1(F74G) enable rapid and extensive inducible degradation, including of very abundant proteins. Considering the generally low basal degradation seen with mIAA7 and OsTIR1(F74G), we chose the mIAA7/a-auxin/OsTIR1(F74G) system for further investigation.

### Impact of OsTIR1(F74G) levels on target protein degradation

The results above indicated that the extent of auxin-induced degradation declines with increasing target protein abundance. A possible reason for this trend is that the levels of the degron receptor limit the capacity of the auxin system. In this case, raising OsTIR1(F74G) levels should enhance the degradation of abundant target proteins. On the other hand, strong OsTIR1(F74G) expression may disturb the ubiquitin-proteasome system because OsTIR1(F74G) competes with other F-box proteins for incorporation into ubiquitin ligase complexes and could thereby interfere with the regular turnover of endogenous proteins. In this case, low OsTIR1(F74G) levels would be desirable. We therefore tested the effect of OsTIR1(F74G) abundance on target protein degradation.

We placed OsTIR1(F74G) under the control of three different promoters, the weak *RNR1* promoter, the strong *ADH* promoter used in all previous experiments, and the extremely strong *GPD* promoter (Figure S5A). Auxin-induced degradation of mIAA7-tagged Fba1, Htb1, Vph1 and Sec63 was faster when OsTIR1(F74G) expression was controlled by the *ADH* rather than the *RNR1* promoter (Figure 4A-D). These results are consistent with a recent study that compared protein degradation by OsTIR1(F74G) expressed under the control of several different constitutive promoters (Fülleborn et al, 2025). However, the differences between the *ADH* and the *RNR1* promoter were minor, and *RNR1*-driven expression was sufficient for almost complete elimination of Sec63. The exceptionally strong *GPD* promoter did not outperform the *ADH* promoter, and in fact gave less complete degradation of Vph1 and Sec63. Hence, OsTIR1(F74G) levels do not limit target protein degradation when the *ADH* promoter is used. The *RNR1* promoter is also a viable choice unless highly abundant proteins are targeted.

**Figure 4.**
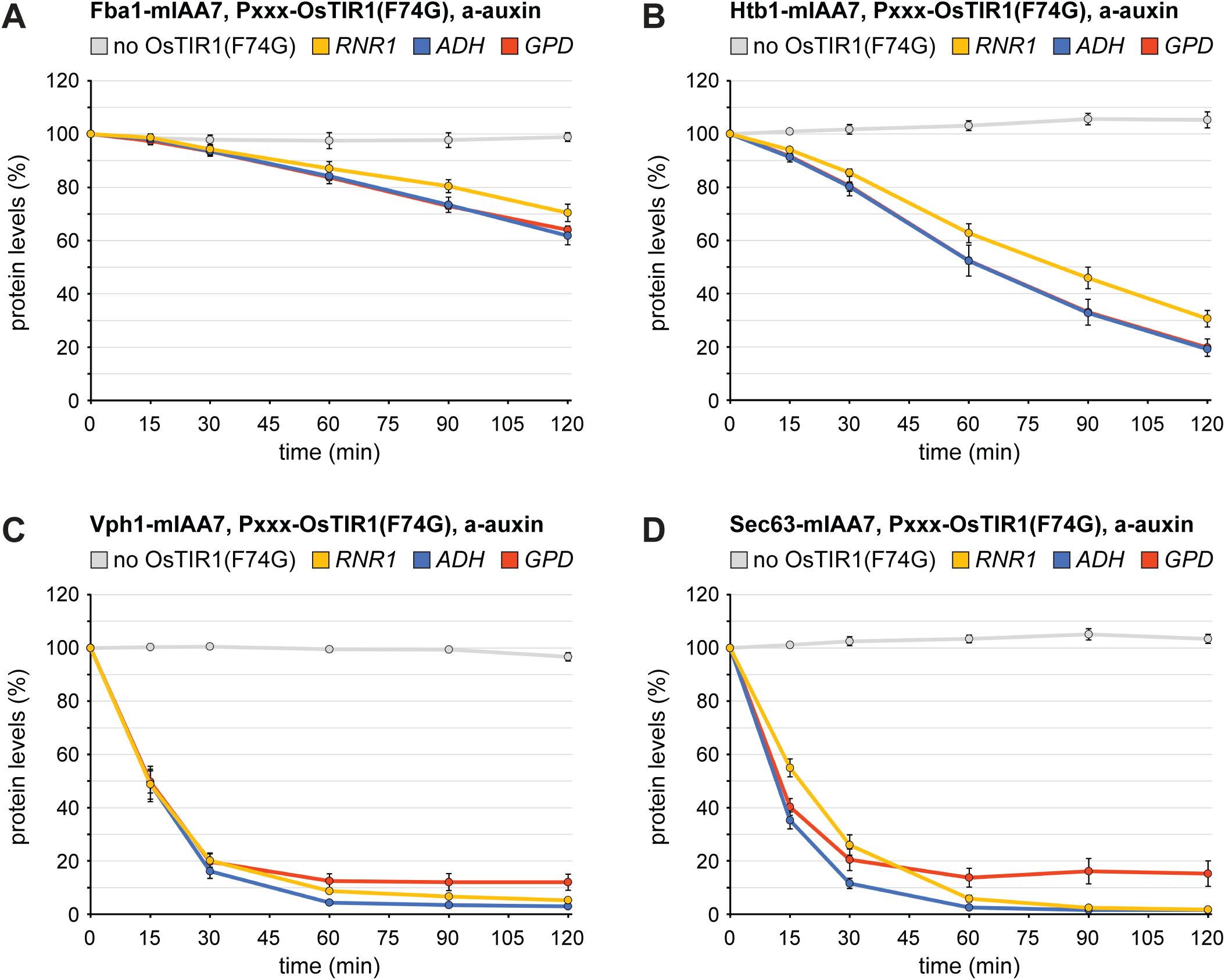
Kinetics of auxin-induced protein degradation by OsTIR1(F74G) expressed via different constitutive promoters. **(A)** Flow cytometry measurement of Fba1-mIAA7- mNeonGreen (Fba1-mIAA7) levels in cells expressing no OsTIR1(F74G) or expressing OsTIR1(F74G) under the control of different constitutive promoters (P_XXX_), i.e. the weak *RNR1* promoter, the strong *ADH* promoter or the very strong *GPD* promoter. Cells were treated with 0.5 µM a-auxin for 0, 15, 30, 60, 90 or 120 minutes. Lines show the mean protein levels of four biological replicates, error bars show the standard error of the mean. For each strain, protein levels were normalised to the levels at t = 0 minutes. **(B-D)** As in panel A but for Htb1, Vph1 and Sec63, respectively.

An alternative way to regulate OsTIR1(F74G) abundance is to use an inducible promoter. This format allows OsTIR1(F74G) to be expressed only at the time of auxin treatment. If the promoter is also titratable, OsTIR1(F74G) levels can be adjusted to the abundance of a target protein. We placed OsTIR1(F74G) under the control of the *GAL* promoter and the artificial transcription factor Gal4-ER-Msn2 (GEM), which activates the *GAL* promoter when the exogenous sterol β-estradiol is added. Furthermore, expression can be adjusted by varying the estradiol concentration (Pincus et al, 2014; Schmidt et al, 2019). To evaluate the potential of this system, we first used it to express superfolder GFP (sfGFP). Baseline sfGFP levels in the absence of estradiol were nearly undetectable, and overnight induction with saturating estradiol concentrations yielded levels similar to those achieved by the *ADH* promoter (Figure S5B). Kinetic measurements showed that sfGFP levels reached those provided by the *RNR1* promoter within one hour of induction and those provided by the *ADH* promoter within three hours (Figure S5C). Hence, the GEM-GAL system shows low baseline expression and allows robust gene induction.

We then combined maximal expression of OsTIR1(F74G) by the GEM-GAL system with auxin-induced protein degradation. Cells in which mIAA7-mNeonGreen was fused to Tef4 or Vph1 and which expressed OsTIR1(F74G) under the control of the *ADH* promoter or the GEM-GAL system were treated with auxin, with estradiol or with both at the same time (Figure 5A, B). In cells constitutively expressing OsTIR1(F74G), auxin reduced Tef4 and Vph1 levels by more than 90% within 80 and 30 minutes, respectively, as observed before (see Figure 3A, C). In cells expressing OsTIR1(F74G) by means of the GEM-GAL system, there was no degradation upon treatment with auxin or estradiol alone, but the combined treatment reduced Tef4 and Vph1 levels by more than 90% within 160 and 90 minutes, respectively. Thus, adding the step of estradiol-induced synthesis of OsTIR1(F74G) delays target protein degradation by roughly one hour. If this delay needs to be avoided, cells can first be treated with estradiol to induce OsTIR1(F74G) expression, and auxin can then be added to trigger target protein degradation.

**Figure 5.**
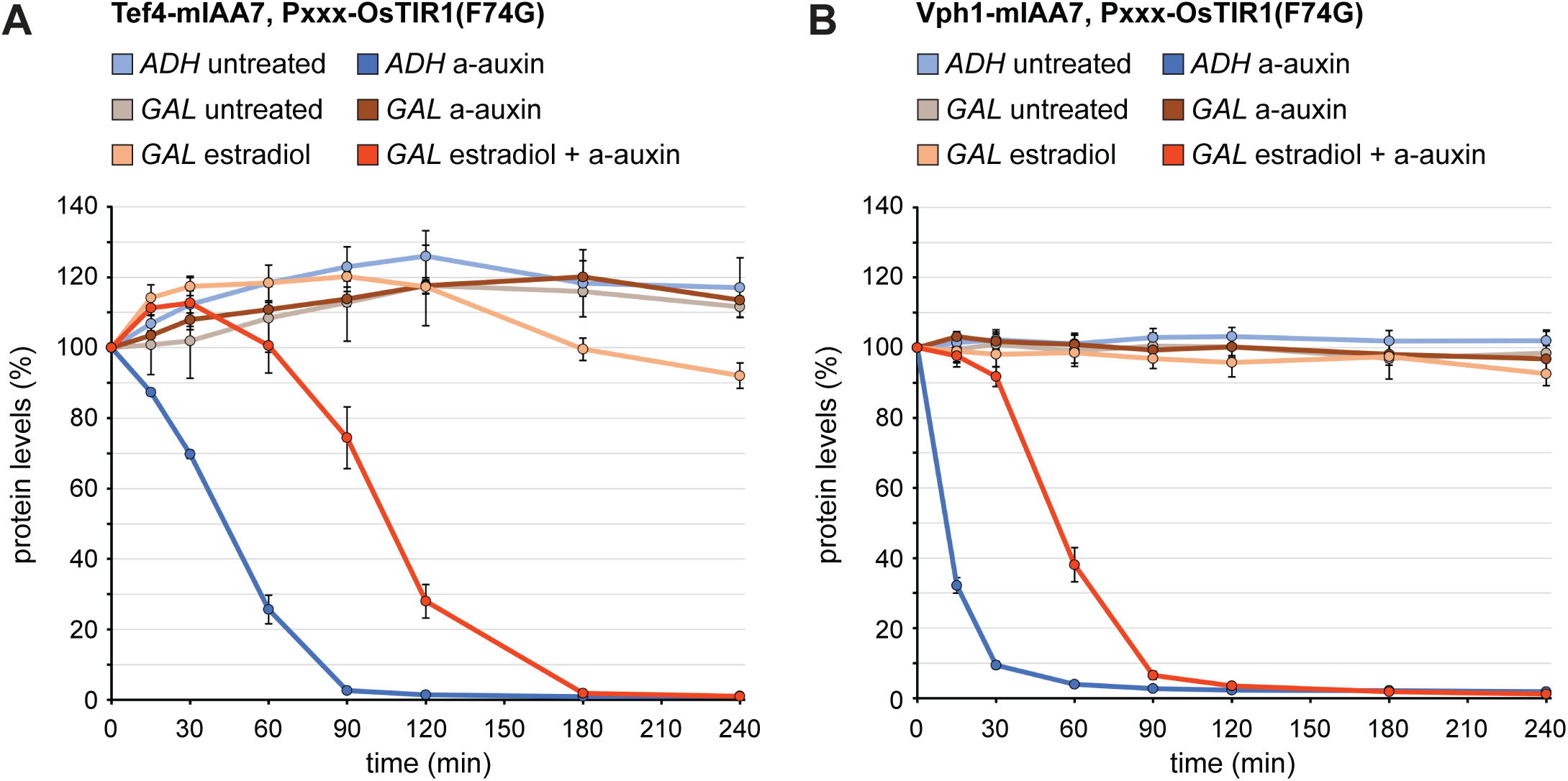
Kinetics of auxin-induced protein degradation by OsTIR1(F74G) expressed via the estradiol-inducible GEM-GAL promoter system. **(A)** Flow cytometry measurement of Tef4-mIAA7-mNeonGreen (Tef4-mIAA7) levels in cells expressing OsTIR1(F74G) under the control of the constitutive *ADH* promoter or the estradiol-inducible GEM-GAL system. Cells were treated with 800 nM estradiol and/or 0.5 µM a-auxin for 0, 15, 30, 60, 90 or 120 minutes. Lines show the mean protein levels of three biological replicates, error bars show the standard error of the mean. For each strain, protein levels were normalised to the levels at t = 0 minutes. **(B)** As in panel A but for Vph1.

In summary, OsTIR1(F74G) expression levels can be adjusted, both constitutively and inducibly, to tune them to target protein abundance. This approach minimises the risk of perturbing the ubiquitin-proteasome system or causing basal target protein degradation.

### Application of the mIAA7/a-auxin/OsTIR1(F74G) system for rapid inhibition of cellular activities

Next, we asked whether rapid protein degradation with the mIAA7/a-auxin/OsTIR1(F74G) system is comparable to protein inactivation with chemical inhibitors and temperature-sensitive (*ts*) alleles. We show that the effects of two widely used chemical inhibitors and a well-characterized *ts* allele can be recapitulated by auxin-inducible depletion of proteins acting in the same cellular pathways.

Tunicamycin is an inhibitor of N-linked protein glycosylation and is commonly used to experimentally induce ER stress. Protein N-glycosylation requires the synthesis of a core oligosaccharide, which is initially anchored in the cytoplasmic leaflet of the ER membrane, then translocated into the ER lumen and finally transferred onto nascent glycoproteins. The first step of core oligosaccharide synthesis is catalysed by Alg7, the molecular target of tunicamycin (Barnes et al, 1984). Alg7 is a multipass transmembrane protein with unknown topology so that it is unclear whether a degron fused to its C-terminus would be exposed to the cytosol. However, the UDP-GlcNAc transferase Alg13, which catalyses the second step of core oligosaccharide synthesis, is a cytosolic protein peripherally associated with the ER membrane (Gao et al, 2005). We therefore fused mIAA7-3xFLAG to the C-terminus of Alg13 to enable its auxin-inducible degradation. Western blotting showed that auxin reduced Alg13 abundance to almost undetectable levels within 15 minutes (Figure 6A). Alg13 is a low-abundance protein (Platzek et al, 2025), which likely explains its very rapid depletion. To assay the consequences of Alg13 degradation, we used a *HAC1* splicing reporter. When N-glycosylation is inhibited, newly synthesised glycoproteins fail to acquire their glycans and cannot fold properly. The resulting accumulation of misfolded proteins in the ER lumen causes ER stress and activates the unfolded protein response (UPR; Walter and Ron, 2011). This signalling pathway is controlled by the ER-resident endoribonuclease Ire1, which, upon ER stress, initiates splicing of the cytosolic *HAC1* mRNA. The *HAC1* splicing reporter is based on the *HAC1* mRNA but translates Ire1 activation into the production of GFP, which can be measured by flow cytometry (Pincus et al, 2010). Tagging of Alg13 with mIAA7-3xFLAG by itself did not cause *HAC1* splicing (Figure 6B, control versus Alg13-mIAA7 strains at the 0-min time point). We then compared activation of the *HAC1* splicing reporter by tunicamycin and auxin-induced Alg13 degradation. Remarkably, Alg13 depletion activated the *HAC1* splicing reporter with similar kinetics and to a similar extent as treatment with 1 µg/ml tunicamycin, which is a standard condition for causing strong ER stress (Figure 6B; Travers et al, 2000; Schuck et al, 2009). Activation of the UPR leads to expansion of the ER (Schuck et al, 2009). Analysis of ER morphology by fluorescence microscopy revealed that Alg13 depletion caused ER expansion, similarly to the effects of 1 µg/ml tunicamycin (Figure 6C). Quantification of ER morphology furthermore showed that expansion was due to a proliferation of ER sheets in both cases (Figure 6D). Hence, Alg13 depletion activates the UPR and induces ER expansion essentially as efficiently as Alg7 inhibition by tunicamycin.

**Figure 6.**
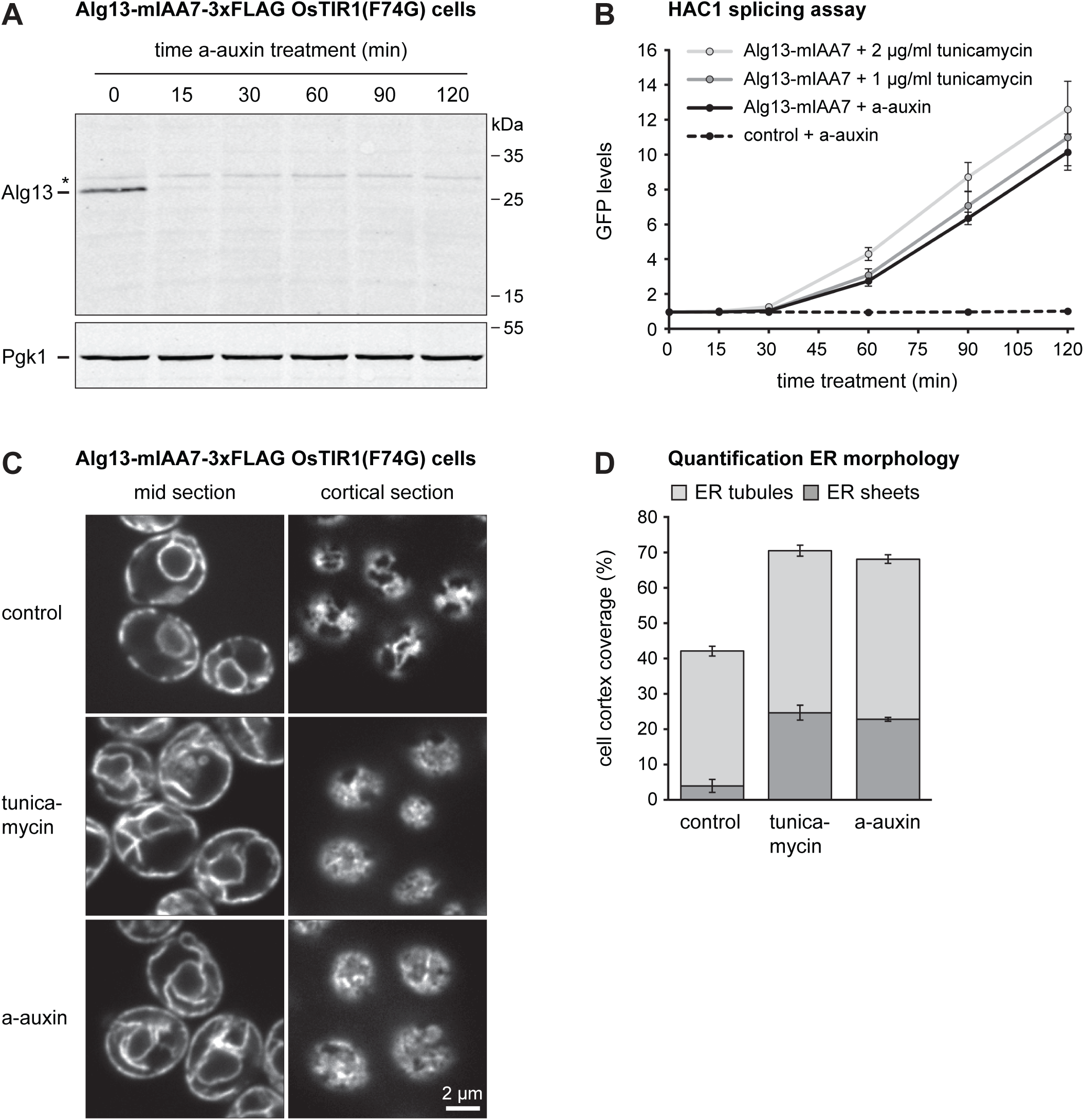
Induction of the unfolded protein response by tunicamycin and auxin-induced Alg13 depletion. **(A)** Western blot of FLAG tag from Alg13-mIAA7-3xFLAG OsTIR1(F74G) cells after treatment with 0.5 µM a-auxin for the times indicated. Pgk1 served as a loading control. The asterisk marks a non-specific band. **(B)** Flow cytometry measurement of GFP produced by the *HAC1* splicing reporter in OsTIR1(F74G) cells (control) and Alg13- mIAA7-3xFLAG OsTIR1(F74G) cells (Alg13-mIAA7) after treatment with 1 µg/ml tunicamycin, 2 µg/ml tunicamycin or 0.5 µM a-auxin. Lines show the mean GFP levels of three biological replicates, error bars show the standard error of the mean. **(C)** Sec63- mNeon images of confocal mid and cortical sections of Sec63-mNeon Alg13-mIAA7- 3xFLAG OsTIR1(F74G) cells that were untreated (control), treated with 1 µg/ml tunicamycin or treated with 0.5 µM a-auxin. Tunicamycin-induced ER stress and auxin- induced Alg13 depletion cause ER expansion. **(D)** Quantification of peripheral ER structures from images as in panel C. Bars are the mean percentage of cell cortex covered by tubules or sheets. Error bars show the 95% confidence interval obtained from at least 12 fields of view per condition, each containing more than 30 cells.

MG132 is a well-established proteasome inhibitor. MG132 has been reported to be ineffective in wild-type yeast but inhibit proteasomal protein degradation in mutants in which net uptake of MG132 into cells has been enhanced. This can be achieved by removal of the ergosterol biosynthesis factor Erg6 or the plasma membrane efflux pump Pdr5 (Lee and Goldberg, 1996; Fleming et al, 2002). Recent high-throughput data demonstrated that the auxin system can be used for proteasomal degradation of many proteasome subunits, i.e. for self-degradation of the proteasome (Gameiro et al, 2025; Valenti et al, 2025). After some initial candidate testing, we settled on Rpn1 as target proteasome subunit. Rpn1 fused to mIAA7 and an ALFA tag (Götzke et al, 2019) was degraded by approximately 75% within 15 minutes of auxin treatment (Figure 7A). Rpn1 levels then remained constant, possibly because a further decline was prevented by a lack of functional proteasomes. To determine whether the reduction of Rpn1 levels was indeed sufficient to reduce proteasome activity, we employed tandem fluorescent timer constructs, called R-tFT and F-tFT (Khmelinskii et al, 2012). These constructs consisted of N-terminal arginine or phenylalanine residues as determinants for proteasomal degradation via the N-degron pathway, a linker sequence that can be ubiquitinated, a slow-maturing mCherry fluorescent protein and a fast-maturing sfGFP fluorescent protein. Newly synthesised reporter molecules rapidly acquire green sfGFP fluorescence and, gradually and more slowly, red mCherry fluorescence. If a reporter is short-lived, it is degraded before maturation of mCherry is complete and therefore shows a low ratio of mCherry to sfGFP fluorescence. Hence, the mCherry/sfGFP fluorescence ratio correlates with protein half-life and provides a readout for reporter stability. We introduced R-tFT and F-tFT reporters into wild-type and *pdr5Δ* cells containing Rpn1-mIAA7-ALFA and OsTIR1(F74G), treated cells with MG132 or a-auxin and measured the mCherry/sfGFP fluorescence ratio by flow cytometry for up to six hours. Treatment with MG132 increased the fluorescence ratio and hence reporter half-life already in wild-type cells and more strongly in *pdr5Δ* cells (Figure 7B, C). For the R-tFT reporter, stabilisation in *pdr5Δ* cells was maximal after two hours and then plateaued. For the F-tFT reporter, stabilisation in *pdr5Δ* cells was clearly apparent after two hours and then further increased up to the six hour time point. Treatment with a-auxin had little effect after two hours but then progressively increased the stabilities of both reporters. After six hours, the increase was comparable to or even somewhat stronger than that caused by MG132. Furthermore, auxin-induced reporter stabilisation was equally strong in wild-type and *pdr5Δ* cells. Hence, the auxin system can be applied in otherwise wild-type cells and achieves similar proteasome inhibition as MG132.

**Figure 7.**
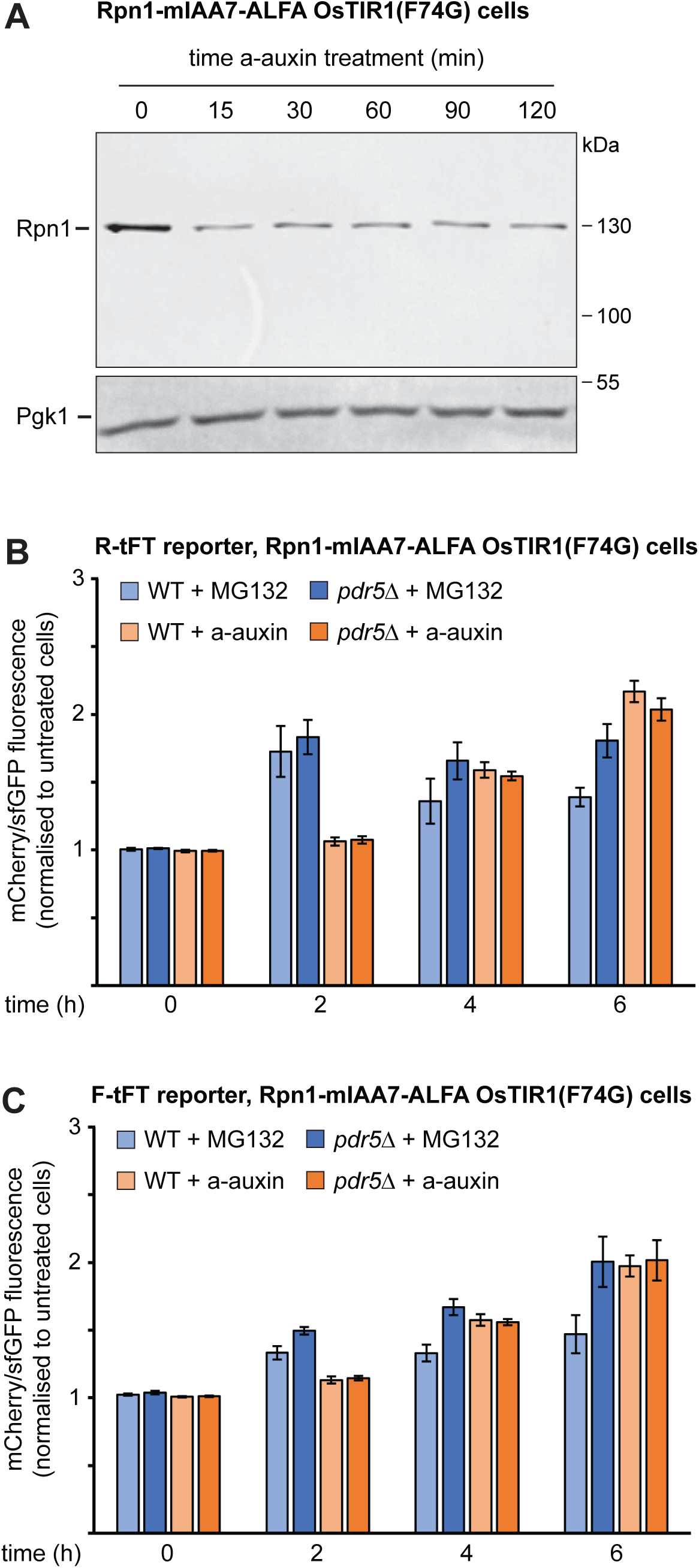
Inhibition of the proteasome by MG132 and auxin-induced Rpn1 depletion. **(A)** Western blot of ALFA tag from Rpn1-mIAA7-ALFA OsTIR1(F74G) cells after treatment with 0.5 µM a-auxin for the times indicated. Pgk1 served as a loading control. **(B)** Flow cytometry measurement of the mCherry/sfGFP fluorescence ratio of the R-tFT reporter in Rpn1-mIAA7-ALFA OsTIR1(F74G) cells that were otherwise wild-type (WT) or lacked the plasma membrane efflux pump Pdr5 (pdr5Δ). Cells were left untreated or treated with 80 µM MG132 or 0.5 µM a-auxin. The mCherry/sfGFP fluorescence ratios in treated cells were normalised to the corresponding ratios in untreated cells. Bars show the mean of four biological replicates, error bars show the standard error of the mean. **(C)** As in panel B but with cells expressing the F-tFT reporter.

Finally, we tested whether the auxin system can be used in place of *ts* alleles. These mutant alleles are important tools to study gene function, especially of essential genes, which cannot be knocked out. Proteins encoded by *ts* alleles are at least partially functional at permissive temperatures but can be rendered non-functional by shifting cells to a different, typically higher, non-permissive temperature. As a test case, we used the established *sec65-1* allele. Sec65 is a subunit of the signal recognition particle and is essential for the co-translational translocation of many secretory proteins into the ER (Hann et al, 1992). When *sec65-1* cells are shifted from the permissive temperature of 25°C to the restrictive temperature of 37°C, signal recognition particle is inactivated and ER insertion of certain newly synthesised proteins fails (Ng et al, 1996). To reproduce this phenotype, we shifted wild-type and *sec65-1* cells from 25°C to 37°C for 60 minutes (Stirling et al, 1992), then induced expression of the vacuolar transmembrane protein GFP-Pho8 with the GEM-GAL system for an additional 60 minutes and finally imaged cells by fluorescence microscopy. As expected, newly synthesised GFP-Pho8 was inserted into the ER and transported to the vacuole membrane under the permissive condition (Figure 8A). By contrast, GFP-Pho8 was unable to reach the vacuole under the restrictive condition and accumulated in cytoplasmic puncta, possibly reflecting aggregation of GFP- Pho8 in the cytosol or even mistargeting to fragmented mitochondria (Costa et al, 2018). In addition, ER structure was disturbed, consistent with earlier observations (Costa et al, 2018). We then generated Sec65-mIAA7-ALFA cells and confirmed by Western blotting that auxin-induced degradation strongly reduced Sec65-mIAA7-ALFA levels within 60 minutes (Figure 8B). Next, we combined auxin-induced degradation of Sec65 for 60 minutes with subsequent estradiol-induced GFP-Pho8 expression for 60 minutes, all at the optimal growth temperature of 30°C. GFP-Pho8 was efficiently transported to the vacuole in the presence of Sec65 but was trapped in cytoplasmic puncta when Sec65 had been depleted. In addition, ER structure was aberrant in Sec65-depleted cells (Figure 8C). Hence, the auxin system presents an alternative to *ts* alleles.

**Figure 8.**
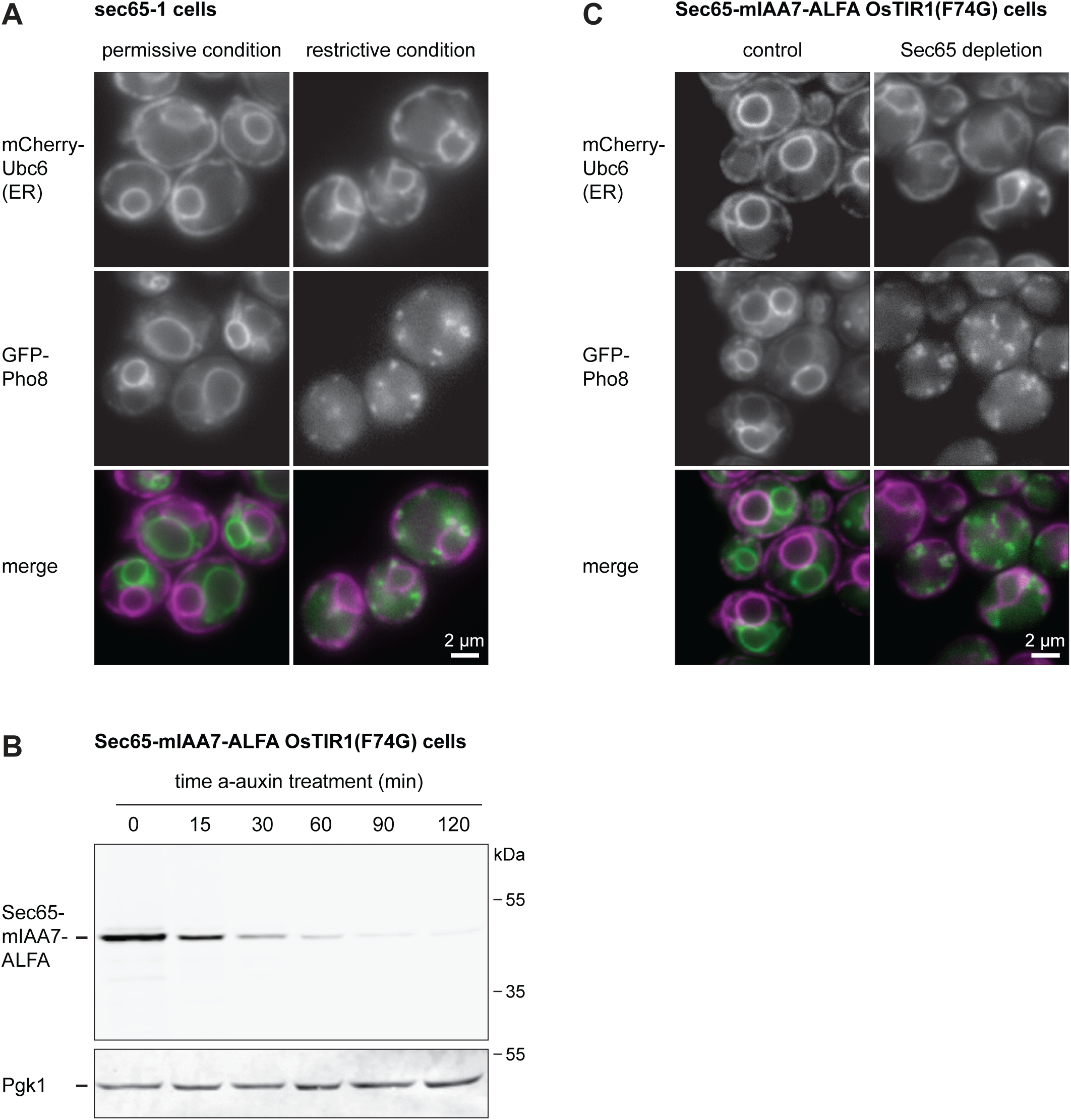
Inactivation of signal recognition particle by the *sec65-1* temperature-sensitive allele or auxin-induced Sec65 depletion. **(A)** Images of *sec65-1* cells constitutively expressing the general ER marker protein mCherry-Ubc6 and expressing the vacuole membrane protein GFP-Pho8 under the control of the estradiol-inducible system. Cells were either grown at 25°C and treated with estradiol at 25°C for 60 min (permissive condition) or grown at 25°C, shifted to 37°C for 60 min and treated with estradiol at 37°C for 60 min (restrictive condition). GFP-Pho8 fails to reach the vacuole membrane under restrictive conditions. **(B)** Western blot of ALFA tag from Sec65-mIAA7-ALFA OsTIR1(F74G) cells after treatment with 0.5 µM a-auxin for the times indicated. Pgk1 served as a loading control. **(C)** Images of Sec65-mIAA7-ALFA OsTIR1(F74G) cells constitutively expressing the general ER marker mCherry-Ubc6 and expressing the vacuole membrane protein GFP-Pho8 under the control of the estradiol-inducible system. Cells were either grown at 30°C and treated with estradiol for 60 min (control) or grown at 30°C, treated with 0.5 µM a-auxin for 60 min and additionally treated with estradiol for another 60 min (Sec65 depletion). Sec65 depletion causes the same Pho8 transport and ER morphology defects observed in *sec65-1* cells at the restrictive temperature.

## Discussion

In this study, we have systematically evaluated combinations of auxin-binding degrons, auxin derivatives and degron receptors. We found that the mIAA7 and AID* degrons together with a-auxin and OsTIR1(F74G) yield rapid and extensive target protein depletion, with little basal degradation in the absence of auxin. We showed that OsTIR1(F74G) should be expressed at high enough but not excessive levels and developed an inducible OsTIR1(F74G) expression system to tune its levels to target protein abundance. This optimised system can replace chemical inhibitors such as tunicamycin or MG132, as well as temperature-sensitive alleles. We have generated plasmids for chromosomal tagging of target proteins with various degron cassettes that include 3xFLAG, ALFA, mNeonGreen or mCherry tags for target protein analysis (Table S3). Furthermore, we have constructed different integrative plasmids for the stable expression of degron receptor variants, including OsTIR1(F74G) under a range of constitutive and inducible promoters (Table S4). All plasmids are available via the European Plasmid Repository and Addgene.

As shown here, and previously by others, the auxin system can target cytosolic proteins, membrane-associated proteins with cytosol-exposed degrons, and also nuclear proteins (Gameiro et al, 2025; Valenti et al, 2025). Hence, OsTIR1(F74G) must have a substantial nuclear pool. Furthermore, the system performs well for transmembrane proteins, which must be extracted from membranes prior to proteasomal degradation, and for proteins that are part of stable complexes, such as histones and proteasomes. The reach of the auxin system is limited for proteins localised entirely inside membrane-bound organelles because these proteins are inaccessible to the proteasome. Still, some depletion may be possible for newly synthesised organelle proteins that are targeted to their final destination post-translationally and can be attacked by the auxin system while they are still in transit. This appears to be the case for a number of mitochondrial proteins (Valenti et al, 2025).

Target protein depletion by about 90% within 30 minutes was achieved for Sec63, Vph1 and Tom70, which are among the 10-20% most abundant proteins in yeast. For proteins among the 5% most abundant proteins, depletion by >90% took longer, as for Tef4 and Htb1, or was not achievable, as for Fba1. We also tested Tdh3, which is the most abundant protein in our W303 strain background (Platzek et al, 2025). Depletion was limited to 10% (unpublished results). This inverse correlation between target protein abundance and depletion efficiency has been noted before (Gameiro et al, 2025; Valenti et al, 2025), but the limiting factor remains unclear. We show that it is not the abundance of the degron receptor. In fact, excessive OsTIR1(F74G) levels can inhibit target protein degradation, as seen for expression driven by the extremely strong *GPD* promoter. An explanation for this counterintuitive effect may be that, at very high concentrations, OsTIR1(F74G) molecules no longer bridge auxin-bound degrons and ubiquitin ligases but separately saturate their respective binding sites at the target protein and the ubiquitin ligase, thus hindering recognition of target proteins by the ubiquitin-proteasome system.

The two degrons that worked well in our hands were AID*, which was recently used for the construction of genome-wide strain collections, and mIAA7 (Murowska and Ulrich, 2013; Li et al, 2019; Gameiro et al, 2025; Valenti et al, 2025). mIAA7 is somewhat larger than AID* (7.5 versus 4.7 kDa) but was slightly superior regarding basal degradation, which is why we used it as our standard degron. We always fused target proteins to cassettes consisting of a degron and a tag so that auxin-induced degradation was easy to monitor. The presence of the additional tag may in some cases interfere with target protein function. However, tags could also stabilise the degron, especially when they contain stably folded protein domains, as is the case for fluorescent proteins. For instance, it has been reported that auxin-inducible degradation of the human seipin protein with mIAA7 alone was slow, was faster when the small 3xFLAG tag was appended to the degron and even faster when a fluorescent protein was used as tag (Li et al, 2019). Hence, it may depend on the target protein whether the best choice is fusion with only a degron, a degron followed by a small epitope tag, or a degron followed by a folded domain. We have not investigated fusions of degrons to the N-termini of target proteins but earlier work indicates that this approach is also possible, including for the mIAA7 degron (Nishimura et al, 2009; Li et al, 2019).

The optimised auxin system is competitive with chemical inhibitors and *ts* alleles. As long as fusion of a degron to a target protein is functionally neutral and a-auxin does not have unintended side effects, the auxin system may in fact offer substantial advantages over inhibitors and *ts* alleles. First, inhibitors can have off-target effects. For instance, tunicamycin causes ER stress but also impacts the mitochondrial proteome in a way that appears unrelated to ER stress (Platzek et al, 2025). Thus, we recommend Alg13 depletion as a new and specific tool to induce ER stress. Second, the auxin system may sometimes be more potent or easier to apply than inhibitors. For instance, it is difficult to introduce MG132 into yeast cells at effective concentrations, even when Erg6 or Pdr5 have been removed or special growth conditions are applied (Lee and Goldberg, 1996; Fleming et al, 2002; Liu et al, 2007). In addition, MG132 only inhibits the chymotryptic activity of the proteasome. Its tryptic and caspase-like activities remain, explaining why MG132 cannot block proteasome activity completely (Collins et al, 2010; Howard et al, 2012). Finally, MG132, which is a peptide aldehyde, appears to be unstable in yeast cells and needs to be replenished after some time (Lee and Goldberg, 1996). Our observation that auxin-induced depletion of Rpn1 in otherwise wild-type cells inhibited the proteasome at least as strongly as MG132 treatment in *pdr5Δ* cells establishes an effective, simple-to- use new tool to study protein degradation in yeast. Rpn1 plays key roles in the delivery of ubiquitinated substrates to the proteasome (Shi et al, 2016). It remains to be determined whether Rpn1 depletion affects all proteasome substrates and whether depletion of another proteasome subunit would provide even tighter proteasome inhibition. Third, *ts* alleles require experimental set-ups in which yeast are shifted from a permissive to a restrictive temperature, neither of which may be optimal for cell physiology. In addition, it needs to be ascertained that any phenotypes result from inactivation of the mutant protein and not from the temperature shift itself. The auxin system can be used at any growth temperature. Fourth, the auxin system requires no target-specific reagents and only involves standard genome manipulation. By contrast, the development of new chemical inhibitors and *ts* alleles cannot be standardised as easily, requires more work, or both.

Considering the efficiency of the auxin system, its application will likely become even more widespread. The reagents now readily available, including genome-wide collections of degron strains (Gameiro et al, 2025; Valenti et al, 2025) and the collection of plasmids described here, make it easy for researchers to adopt the auxin system as a standard approach to investigate protein function in yeast. We expect that the consistent use of auxin-inducible protein degradation will alleviate problems caused by cell adaptation and off-target effects and enable new biological insight.

## Materials and Methods

### Plasmids

Plasmids used in this study are listed in Table S1. Plasmids generated in this study as tools for the auxin system (Tables S3 and S4) are available from the European Plasmid Repository (www.plasmids.eu) and Addgene (www.addgene.org).

Plasmids of the pFA6a series have been described (Longtine et al, 1998, Janke et al, 2004; Papagiannidis et al, 2021). The new kanNP1 module consists of the *S. cerevisiae TPI1* promoter, the *E. coli* transposon *Tn903* kanR open reading frame and the *A. gossypii TEF* terminator. It replaces the kanMX6 module and avoids unwanted recombination in yeast containing both the kanMX6 and the HIS3MX6 module. mAID, AID* and mIAA7 sequences (Yesbolatova et al, 2020; Morawska and Ulrich, 2013; Li et al, 2019) were inserted into pFA6a plasmids as synthetic gene fragments by Gibson assembly cloning. The OsTIR1(F74G) sequence was amplified from P_GAL_-OsTIR1(F74G)-URA3 (Yesbolatova et al, 2020) and inserted into pRS405/6-P_ADH_ (Mumberg et al, 1995). The P_ADH_-OsTIR1(F74G) cassette was then inserted into pNH604/5 vector backbones (Pincus et al, 2014). In the process, the *ADH* promoter was shortened to approximately 700 bp and thus slightly weakened, as in earlier pNH604/5 plasmids (Pincus et al, 2014). Similarly, the AtAFB2 sequence from pSH-EFIRES-B-AtAFB2-mCherry (Li et al, 2019) was inserted into pRS405-P_ADH_ and the P_ADH_-AtAFB2 cassette with the shortened *ADH* promoter was then inserted into the pNH605 vector backbone. Site-directed mutagenesis was used to generate the variants OsTIR1, OsTIR1(F74A), AtAFB2(F74A) and AtAFB2(F74G). To generate pRS303K-P_GPD_-sfGFP, pRS303K-P_GPD_-TagBFP (Szoradi et al, 2018) was linearised by PCR and combined with sfGFP sequence by Gibson assembly cloning, replacing TagBFP. To generate pRS303K-P_RNR1_-sfGFP and pRS303K-P_ADH_- sfGFP, the *RNR1* and the shortened *ADH* promoter were amplified from pYTK021 (Lee et al, 2015) and pNH604-P_ADH_-OsTIR1(F74G), respectively, and inserted into pRS303K- P_GPD_-sfGFP, replacing the *GPD* promoter. To generate pRS303K-P_XXX_-OsTIR1(F74G) plasmids, the OsTIR1(F74G) sequence was amplified from pNH604-P_ADH_-OsTIR1(F74G) and inserted into pRS303K-P_GPD_-sfGFP, pRS303K-P_ADH_-sfGFP and pRS303K-P_RNR1_- sfGFP, replacing sfGFP. pRS303H/N-OsTIR1(F74G) were generated by insertion of OsTIR1(F74G) into pRS303H/N (Taxis and Knop, 2006). pRS303K-P_GAL_-sfGFP was generated by replacing the *RNR1* promoter in pRS303K-P_RNR1_-sfGFP with the *GAL* promoter from pNH605-P_ADH_-GEM-P_GAL_ (Schmidt et al, 2019). Next, the P_ADH_-GAL4-ER- Msn2 (GEM) sequence from pNH605-P_ADH_-GEM-P_GAL_ was inserted, yielding pRS303K- P_GAL_-sfGFP-P_ADH_-GEM. Finally, sfGFP was replaced by the OsTIR1(F74G) sequence, yielding pRS303K-P_GAL_-OsTIR1(F74G)-P_ADH_-GEM.

### Yeast strains

Strains in this study were in the W303 background (genotype *ADE2 leu2-3,112 trp1-1 ura3-1 his3-11,15 MATa*) and are listed in Table S2. Gene tagging and deletion was done with PCR products (Longtine et al, 1998; Janke et al, 2004). Table S3 lists the primers to be used with each template plasmid for gene tagging. For a given gene, primers R1 and S2 can be used interchangeably but primers F2 and S3 cannot. Even though all pFA6a plasmids have sequences complementary to both the F2 and S3 primers, each pFA6a plasmid yields a PCR product with the correct reading frame with only one of them. Expression plasmids for the degron receptors were digested with restriction enzymes prior to integration into the genome. Table S4 lists the restriction enzymes that can be used for this purpose. Note that multiple copies of linearised pRS405/6 plasmids can integrate into the genome, whereas pNH604/5 and pRS303H/K/N are single-integration plasmids.

### Flow cytometry

Strains were grown in liquid culture at 30°C in synthetic complete (SC) medium containing 0.7% yeast nitrogen base without amino acids (Merck), amino acids and 2% glucose. Precultures were inoculated in 1 ml medium in 96-deep well plates and grown for 24 h so that they reached saturation. Precultures were diluted into 1 ml fresh medium and grown overnight so that they reached mid log phase (OD_600_ = 0.1 – 0.5). Overnight cultures were diluted to OD_600_ = 0.05 and then either not treated or treated with 375 µM the natural 3- indoleacetic acid (n-auxin, Sigma-Aldrich), 5 µM phenyl-3-indoleacetic acid (p-auxin, Sigma-Aldrich), 0.5 µM adamantyl-3-indoleacetic acid (a-auxin, TCI Chemicals), 800 nM estradiol (Sigma-Aldrich), 1 or 2 µg/ml tunicamycin (Sigma-Aldrich) or 80 µM MG132 (Thermo Fisher Scientific) for the times indicated. Total cell fluorescence after excitation with a 488 nm laser was measured with a FACSCanto or a FACSymphony A1 flow cytometer (BD Biosciences), and the geometric mean of the cell population was calculated with FlowJo software. Cell autofluorescence was determined with a strain not expressing a fluorescent protein and subtracted from all other measurements. Cellular protein levels were determined by dividing the background-corrected cell fluorescence by the forward scatter, which served as a measure for cell size.

For the analysis of UPR activation by means of the *HAC1* splicing reporter, cells were used that expressed cytosolic BFP as an indicator for cellular protein translation capacity. Background-corrected GFP fluorescence was divided by background-corrected BFP fluorescence elicited with a 405 nm laser, and the GFP/BFP ratio in treated cells was normalised to the ratio in untreated cells, yielding fold induction of the reporter upon tunicamycin or auxin treatment.

For analysis of proteasome activity, mCherry and sfGFP fluorescence was measured in cells expressing R-tFT or F-tFT reporters after excitation with a 561 nm and a 488 nm laser, respectively, values were corrected for autofluorescence by subtracting values for mCherry and sfGFP fluorescence determined in cells not expressing a tFT reporter, and the ratio of background-corrected mCherry/sfGFP fluorescence was calculated. The mCherry/sfGFP ratio in treated cells was normalised to the ratio in untreated cells.

### Western blotting

Strains were grown to mid log phase in liquid culture at 30°C in SC medium. Cultures were diluted to OD_600_ = 0.3 and treated with 0.5 µM a-auxin for up to two hours. At each time point, five ODs of cells were collected by centrifugation, washed with water, resuspended in lysis buffer (50 mM HEPES pH 7.5, 0.5 mM EDTA, Roche complete protease inhibitors) and disrupted by bead beating with a FastPrep 24 (MP Biomedicals). Proteins were solubilised by addition of 1.5% (w/v) SDS and incubation at 65°C for 5 min. Lysates were cleared at 16,000 g at 4°C for 2 min and protein concentrations were determined with a BCA kit (Pierce). Twenty micrograms total cell protein were separated by Tris-glycine SDS-PAGE and transferred onto PVDF membranes by semi-dry blotting. Membranes were probed with primary and fluorophore-coupled secondary antibodies. For fluorescence detection, membranes were analysed with an Odyssey CLx imaging system and Image Studio software (LI-COR Biosciences). Antibodies were mouse anti-FLAG (clone M2, Merck, RRID:AB_262044), mouse anti-Pgk1 (clone 22C5D8, Abcam, RRID:AB_10861977), single-domain anti-ALFA coupled to IRDye 800CW (clone 1G5, NanoTag Biotechnologies, RRID:AB_3075985), goat anti-mouse IRDye 800CW (LI-COR, RRID:AB_621842) and goat anti-mouse Alexa-680 (Invitrogen, RRID:AB_141436).

### Microscopy

To image ER remodelling upon ER stress or Alg13 depletion, strains were grown to mid log phase in liquid SC medium at 30°C. Cultures were diluted to OD_600_ = 0.2 and either not treated or treated with 1 µg/ml tunicamycin or 0.5 µM a-auxin for 2 hours. Cells from 1 ml of culture were collected by centrifugation and resuspended in 20 µl SC. 3 µl cell suspension were applied to a glass coverslip and covered with a 1% agarose pad prepared with SC medium. Images were acquired with a DMi8 inverted microscope (Leica) equipped with a CSU-X1 spinning-disk confocal scanning unit (Yokogawa) and an ORCA- Flash 4.0 LT camera (Hamamatsu). A HC PL APO 63x/1.40-0.60 oil objective lens (Leica) was used. For image analysis, images were anonymised with the “Blind Analysis Tools” plugin in ImageJ (https://imagej.net/plugins/blind-analysis-tools) to prevent user bias. Cells were then segmented using CellX software (Dimopoulos et al, 2014), and ER morphology (% cell cortex coverage, ER tubules and ER sheets) was quantified with the ClassifiER script in MATLAB (Papagiannidis et al, 2021). At least 12 fields of view with at least 34 cells each were quantified per condition, and a 95% confidence interval was calculated based on the variance between fields of view.

To image ER translocation of newly synthesised GFP-Pho8, strains were grown to mid log phase in liquid SC medium at 25°C (*sec65-1* strain) or 30°C (Sec65-mIAA7-ALFA strain). Cultures of *sec65-1* cells were split in two, maintained at 25°C or shifted to 37°C for 60 min, treated with 800 nM estradiol to induced GFP-Pho8 expression for 60 min and imaged with a Nikon Ti2 widefield microscope equipped with a Nikon Plan Apo 100x/NA 1.45 objective and a Hamamatsu Orca Fusion-BT camera. Cultures of Sec65-mIAA7- ALFA cells were split in two, not treated or treated with 0.5 µM a-auxin for 60 min, and then treated with estradiol and imaged as above.

## Acknowledgements

We thank the Flow Cytometry & FACS Core Facility at the Center for Molecular Biology at Heidelberg University (ZMBH) for assistance, Heike Adler for help with plasmid generation, Michael Knop for plasmids and all Schookees for comments on the manuscript. This project was supported, in part, by grant SCHU 2364/3-1 from the German Research Foundation (DFG) to SS. The authors gratefully acknowledge the data storage service SDS@hd supported by the Ministry of Science, Research and the Arts Baden- Württemberg (MWK) and the DFG through grant INST 35/1503-1 FUGG. The authors declare no competing financial interests.

## Author contributions

Conceptualisation: Nadja Guschtschin-Schmidt, Petra Hubbe, Oliver Pajonk, Sebastian Schuck; Investigation: Natalie Friemel, Nadja Guschtschin-Schmidt, Petra Hubbe, Oliver Pajonk, Niklas Peters, Sebastian Schuck, Charu Sharma; Supervision and Writing – original draft: Sebastian Schuck; Writing – review and editing: all authors.

---

**Figure S1.**
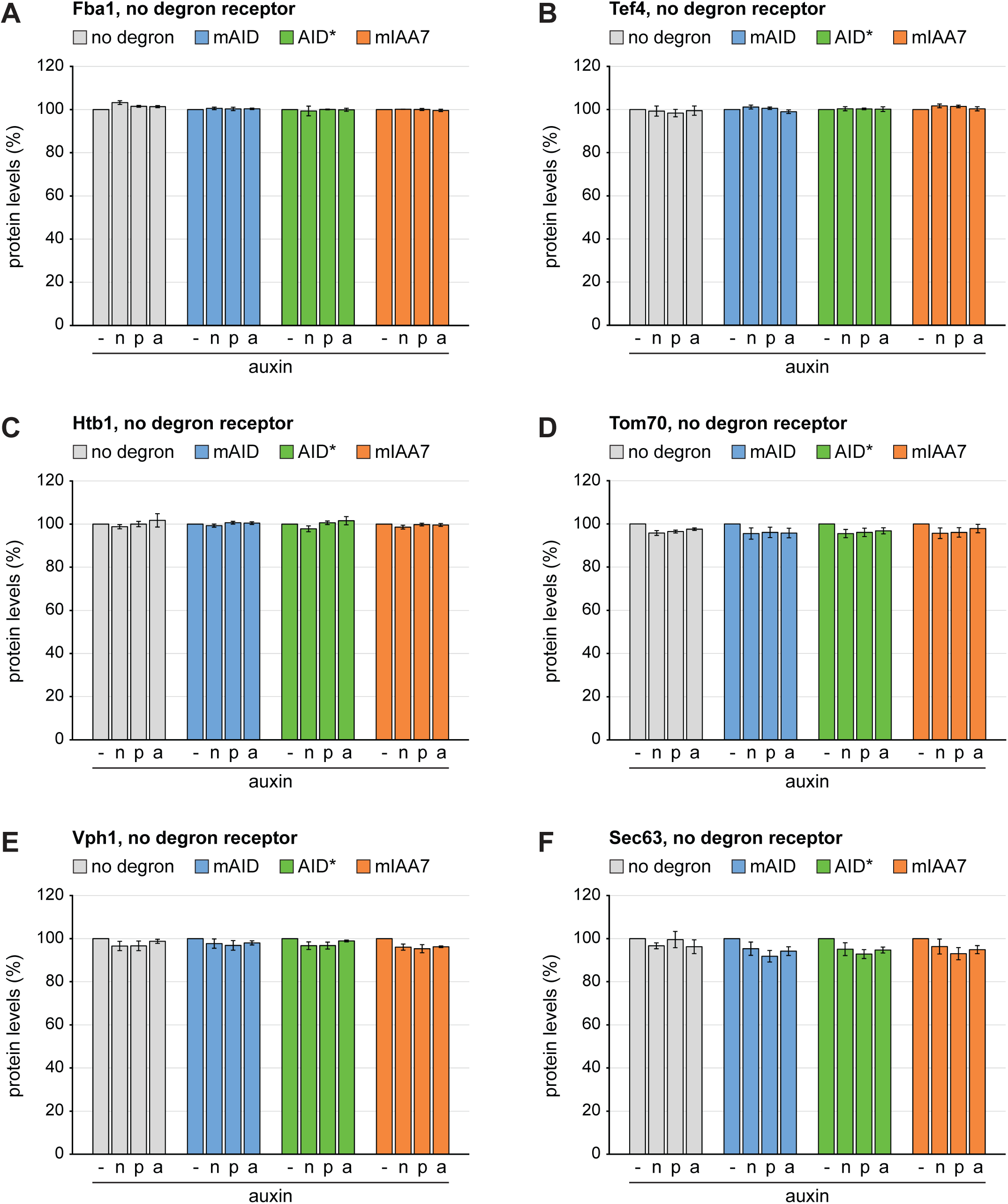
Basal degradation by auxin in the absence of a degron receptor. **(A)** Flow cytometry measurement of relative levels of Fba1 tagged with mNeonGreen (no degron), mAID-mNeonGreen (mAID), AID*-mNeonGreen (AID*) or mIAA7-mNeonGreen (mIAA7). Cells did not express a degron receptor and were not treated or treated with 375 µM n- auxin, 5 µM p-auxin or 0.5 µM a-auxin for 120 minutes. Bars show the mean protein levels of at least three biological replicates, error bars show the standard error of the mean. Protein levels were normalised to the levels in untreated cells. **(B-F)** As in panel A but for Tef4, Htb1, Tom70, Vph1 and Sec63, respectively.

**Figure S2.**
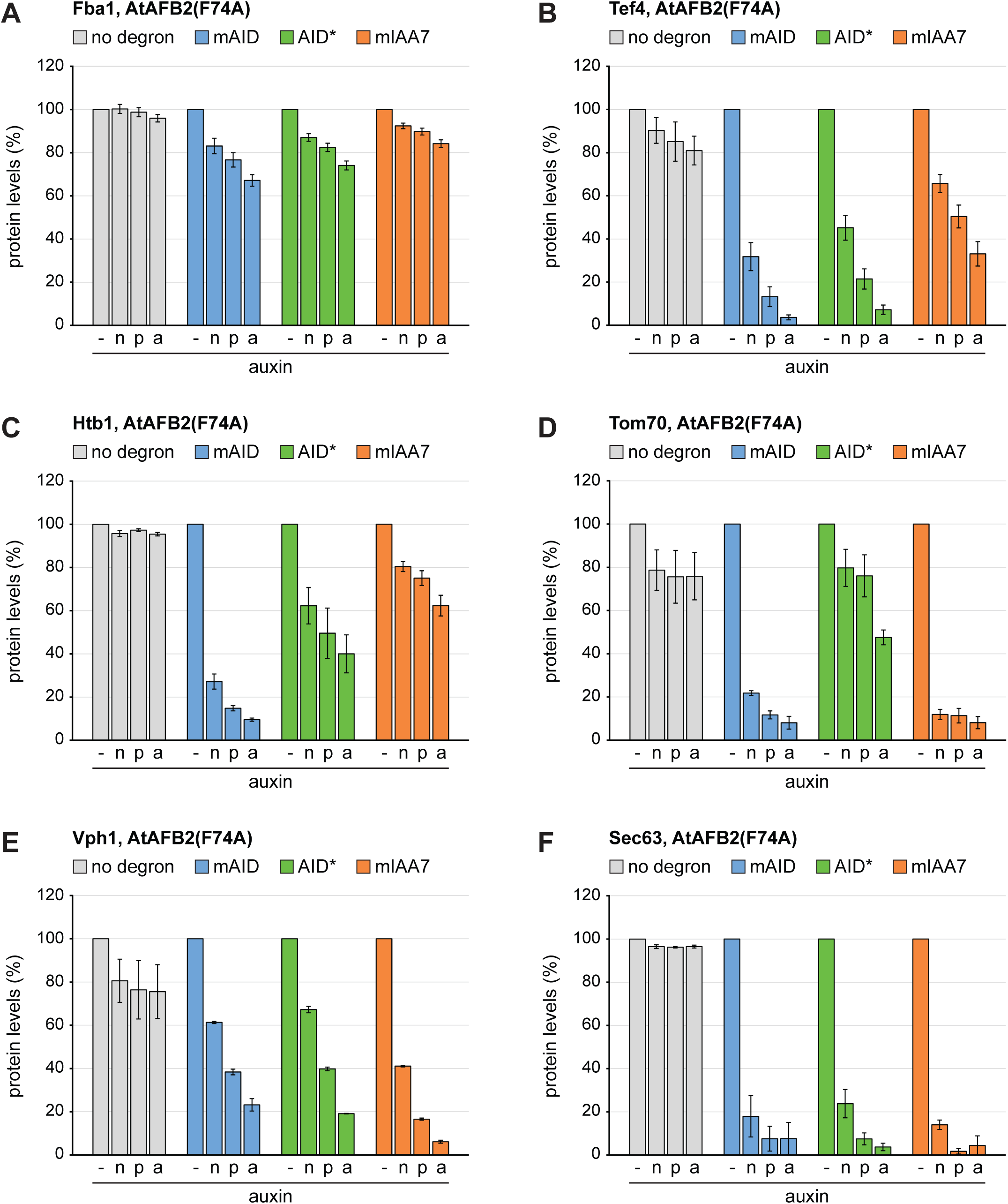
Inducible protein degradation by n-/p-/a-auxin in the presence of AtAFB2(F74A). **(A)** Flow cytometry measurement of relative levels of Fba1 tagged with mNeonGreen (no degron), mAID-mNeonGreen (mAID), AID*-mNeonGreen (AID*) or mIAA7-mNeonGreen (mIAA7). Cells expressed AtAFB2(F74A) and were not treated or treated with 375 µM n-auxin, 5 µM p-auxin or 0.5 µM a-auxin for 120 minutes. Bars show the mean protein levels of three biological replicates, error bars show the standard error of the mean. Protein levels were normalised to the levels in untreated cells. **(B-F)** As in panel A but for Tef4, Htb1, Tom70, Vph1 and Sec63, respectively.

**Figure S3.**
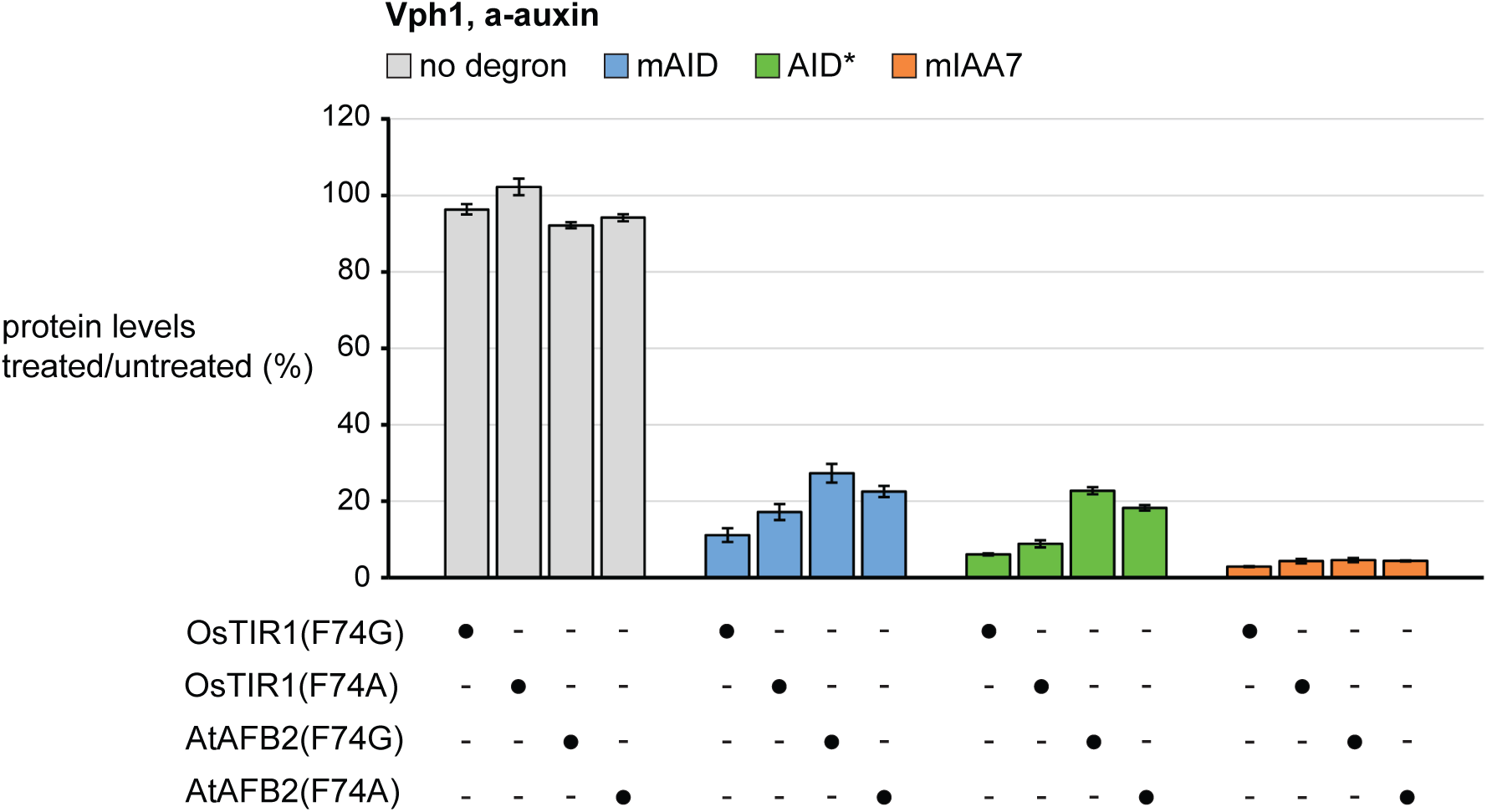
Auxin-inducible degradation by different degron receptor variants. Flow cytometry measurement of relative levels of Vph1 tagged with mNeonGreen (no degron), mAID-mNeonGreen (mAID), AID*-mNeonGreen (AID*) or mIAA7-mNeonGreen (mIAA7). Cells expressed OsTIR1(F74G), OsTIR1(F74A), AtAFB2(F74G) or AtAFB2(F74A) and were not treated or treated 0.5 µM a-auxin for 120 minutes. Bars show the mean protein levels in treated relative to untreated cells of three biological replicates, error bars show the standard error of the mean.

**Figure S4.**
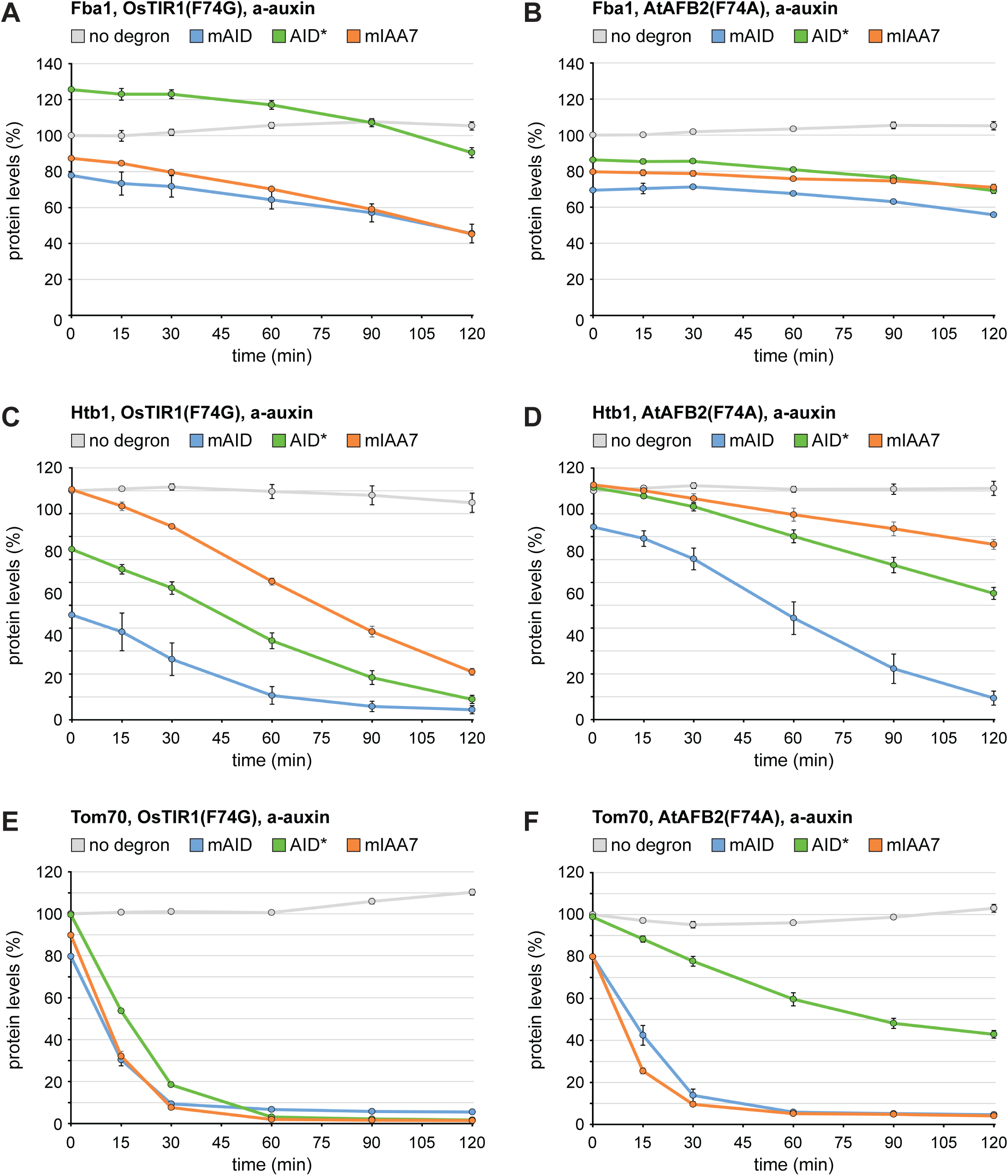
Kinetics of inducible protein degradation by a-auxin in the presence of OsTIR1(F74G) or AtAFB2(F74A). **(A)** Flow cytometry measurement of relative levels of Fba1 tagged with mNeonGreen (no degron), mAID-mNeonGreen (mAID), AID*- mNeonGreen (AID*) or mIAA7-mNeonGreen (mIAA7). Cells expressed OsTIR1(F74G) and were treated with 0.5 µM a-auxin for 0, 15, 30, 60, 90 or 120 minutes. Lines show the mean protein levels of three biological replicates, error bars show the standard error of the mean. Protein levels were normalised to the levels in cells that expressed Tef4- mNeonGreen (no degron) and were treated with a-auxin for 0 min. **(B)** As in panel A but for cells expressing AtAFB2(F74A). **(C, E)** As in panel A but for cells expressing tagged variants of Htb1 and Tom70, respectively. **(D, F)** As in panel A but for cells expressing AtAFB2(F74A) and tagged variants of Htb1 and Tom70, respectively.

**Figure S5.**
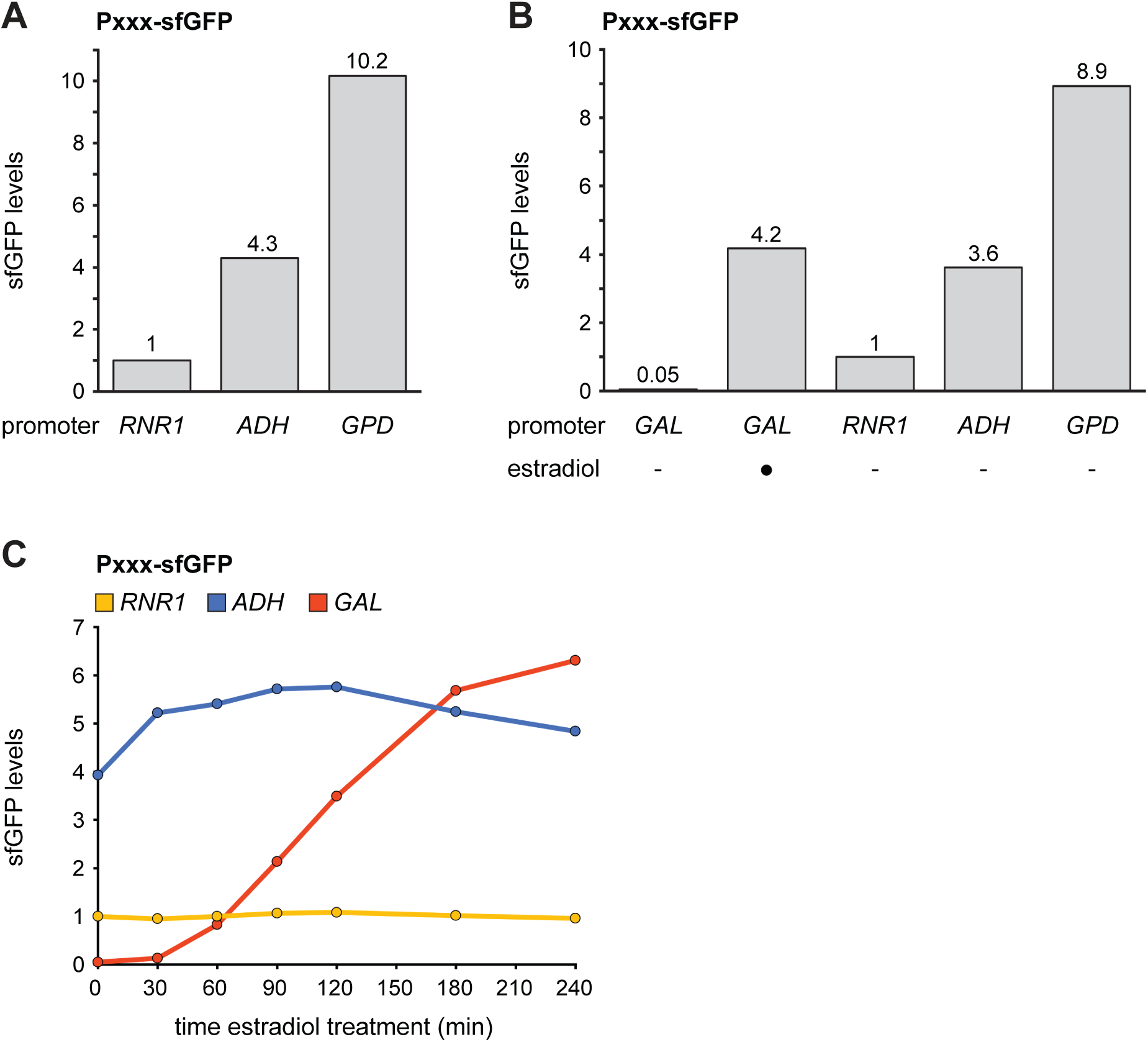
Strength of constitutive and inducible promoters used for OsTIR1(F74G) expression. **(A)** Flow cytometry measurement of sfGFP levels in cells expressing sfGFP under the control of different constitutive promoters (P_XXX_), i.e. the *RNR1*, *ADH* or *GPD* promoter. Bars show the mean of two biological replicates. sfGFP levels were normalised to the levels in cells expressing sfGFP under the control of the *RNR1* promoter. **(B)** As in panel A but including the GEM-GAL promoter system. Cells were grown in the absence of estradiol or treated with 800 nM estradiol for 16 hours. Bars show the mean of two technical replicates. sfGFP levels were normalised to the levels in cells expressing sfGFP under the control of the *RNR1* promoter. **(C)** As in panel A but showing sfGFP levels over time after treatment with 800 nM estradiol for up to four hours. Lines show the mean of two technical replicates. sfGFP levels were normalised to the levels in cells expressing sfGFP under the control of the *RNR1* promoter at t = 0 minutes.

**Table S1.**
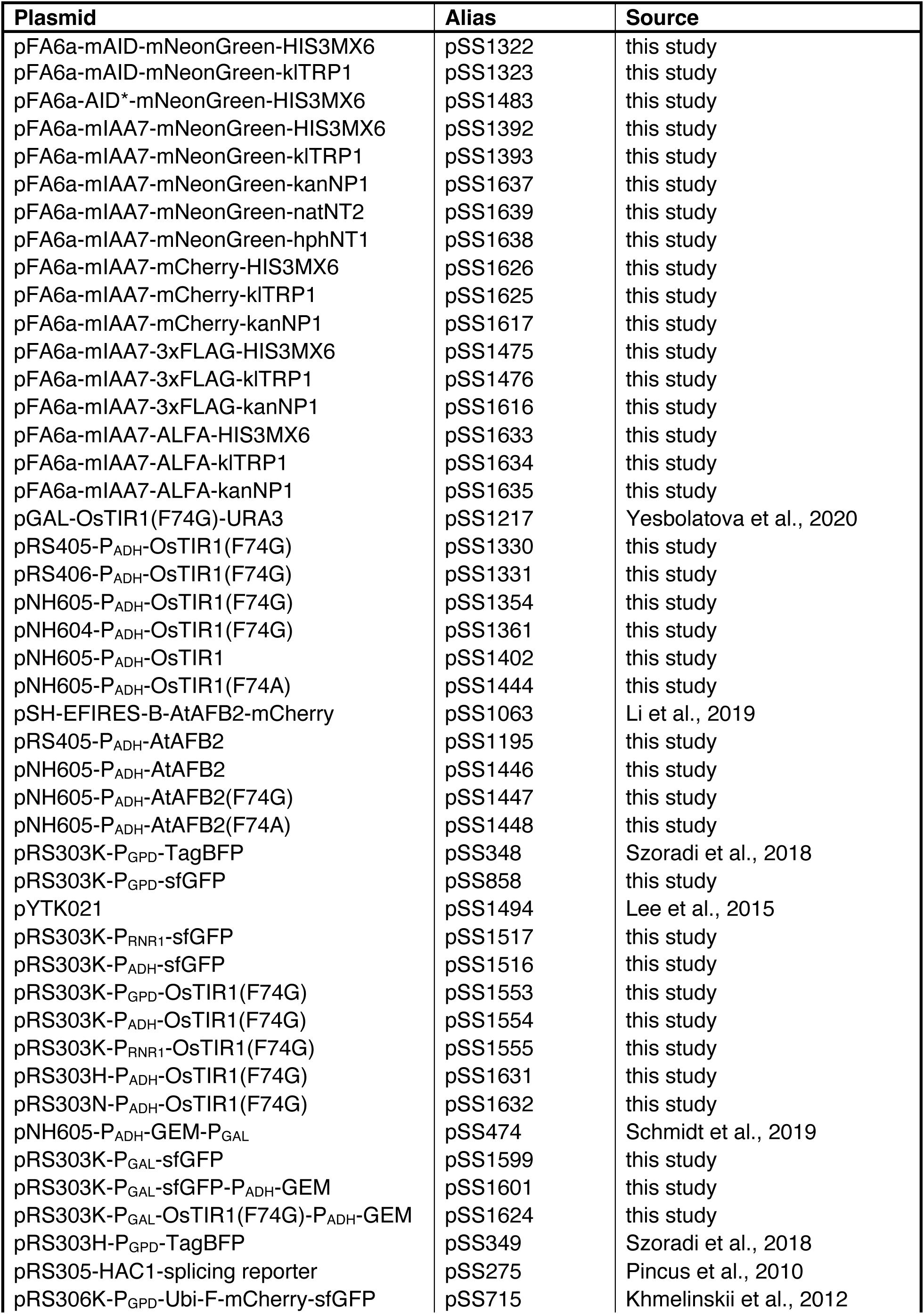
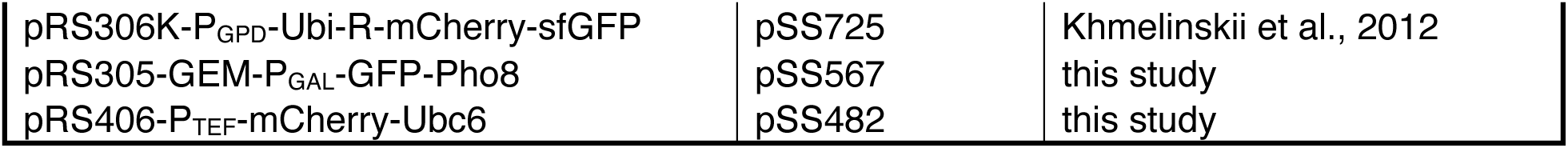
Plasmids used in this study. GEM, GAL4-ER-Msn2; sfGFP, superfolder GFP.

**Table S2.**
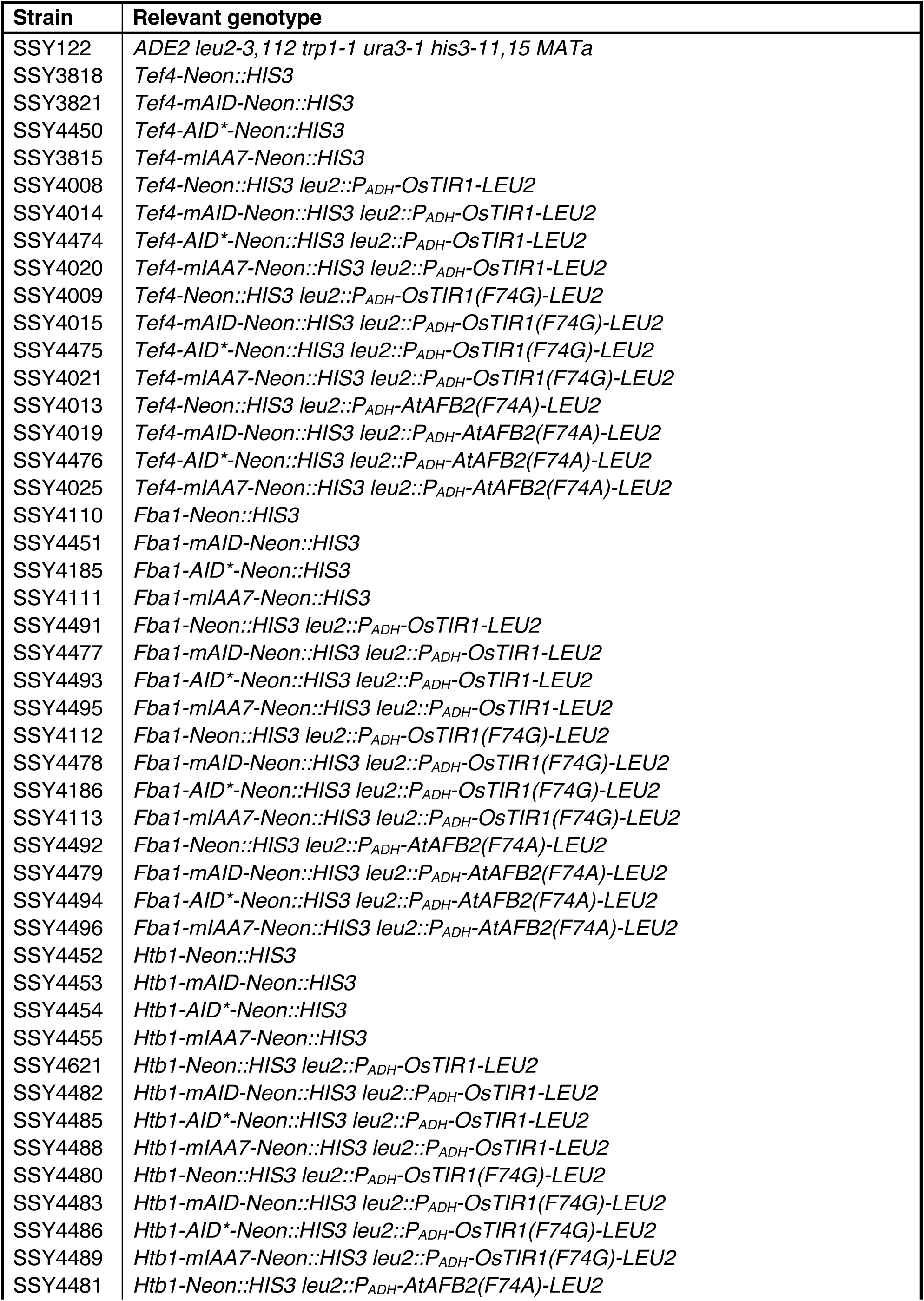
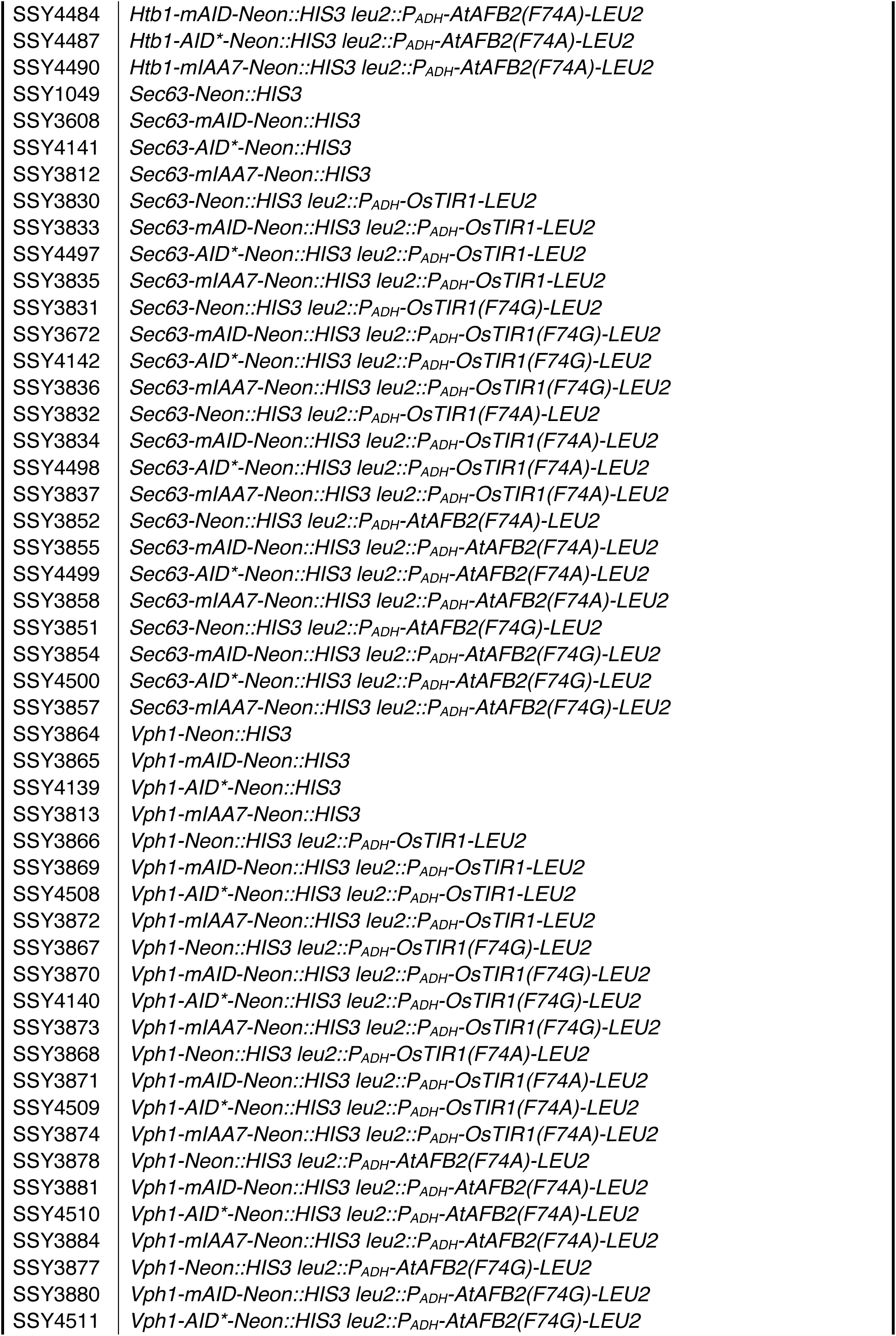
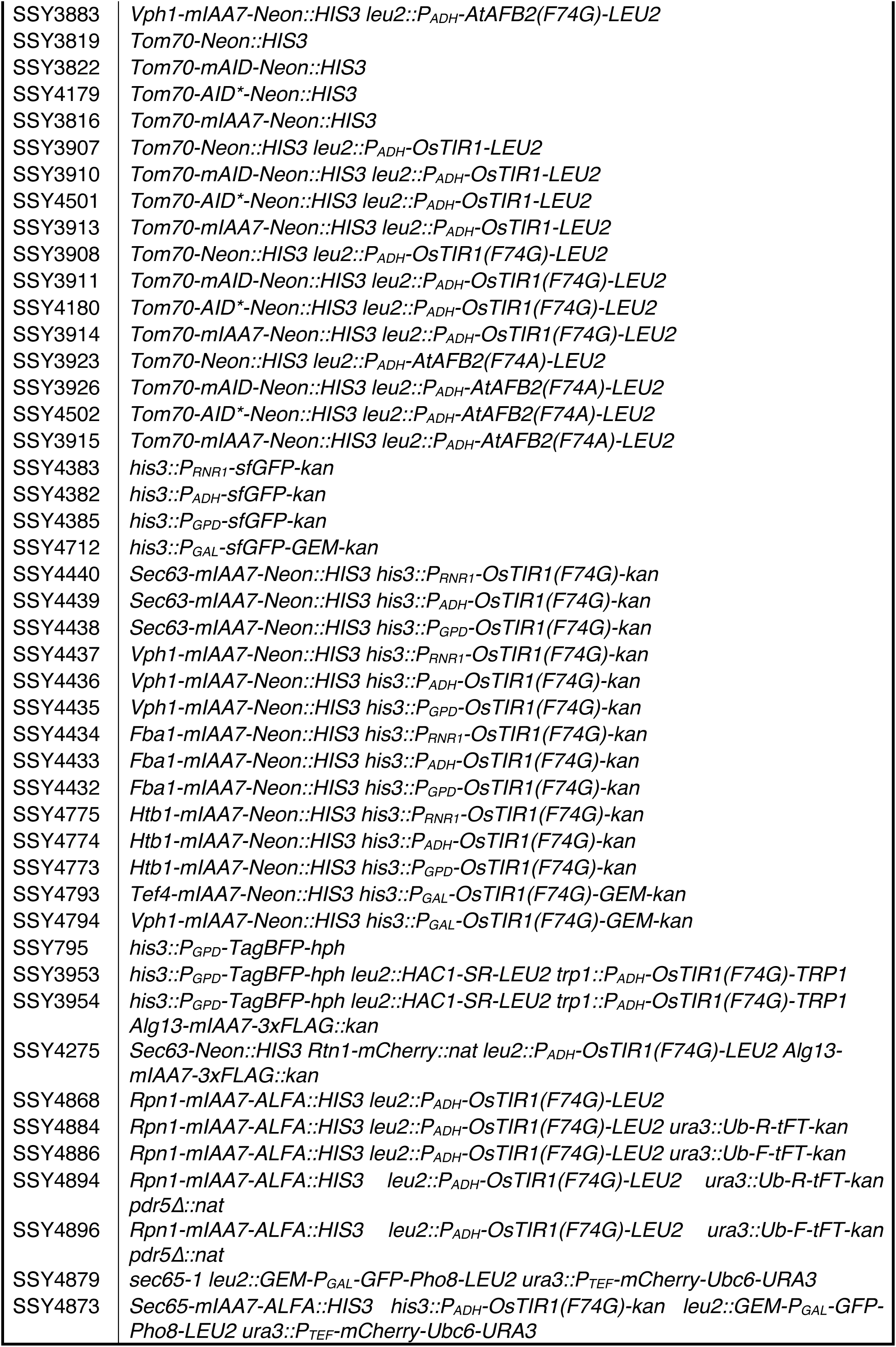
Yeast strains used in this study. GEM, GAL4-ER-Msn2; Neon, mNeonGreen; sfGFP, superfolder GFP; SR, splicing reporter.

**Table S3.** Plasmids and cognate forward and reverse primers for gene tagging. F2 primers consist of the 40-nucleotide sequence immediately upstream of the stop codon of a target gene (excluding the stop codon), followed by 5’-CGGATCCCCGGGTTAATTAA- 3’. R1 primers consist of the reverse complement of the 40-nucleotide sequence immediately downstream of the stop codon of a target gene (excluding the stop codon), followed by 5’-GAATTCGAGCTCGTTTAAAC-3’ (Longtine et al, 1998). S3 primers consist of the 42-nucleotide sequence immediately upstream of the stop codon of a target gene (excluding the stop codon), followed by 5’-CGTACGCTGCAGGTCGAC-3’. S2 primers consist of the reverse complement of the 41-nucleotide sequence immediately downstream of the stop codon of a target gene (excluding the stop codon), followed by 5’- ATCGATGAATTCGAGCTCG-3’ (Janke et al, 2004).

| Plasmid | Alias | forward primer | reverse primer |
| --- | --- | --- | --- |
| pFA6a-mAID-Neon-HIS3MX6 | pSS1322 | F2 | R1 or S2 |
| pFA6a-mAID-Neon-kITRP1 | pSS1323 | F2 | R1 or S2 |
| pFA6a-AID*-Neon-HIS3MX6 | pSS1483 | F2 | R1 or S2 |
| pFA6a-mIAA7-Neon-HIS3MX6 | pSS1392 | F2 | R1 or S2 |
| pFA6a-mIAA7-Neon-kITRP1 | pSS1393 | F2 | R1 or S2 |
| pFA6a-mIAA7-Neon-kanNP1 | pSS1637 | F2 | R1 or S2 |
| pFA6a-mIAA7-Neon-natNT2 | pSS1639 | S3 | R1 or S2 |
| pFA6a-mIAA7-Neon-hphNT1 | pSS1638 | S3 | R1 or S2 |
| pFA6a-mIAA7-Cherry-HIS3MX6 | pSS1626 | F2 | R1 or S2 |
| pFA6a-mIAA7-Cherry-kITRP1 | pSS1625 | F2 | R1 or S2 |
| pFA6a-mIAA7-Cherry-kanNP1 | pSS1617 | F2 | R1 or S2 |
| pFA6a-mIAA7-3xFLAG-HIS3MX6 | pSS1475 | F2 | R1 or S2 |
| pFA6a-mIAA7-3xFLAG-kITRP1 | pSS1476 | F2 | R1 or S2 |
| pFA6a-mIAA7-3xFLAG-kanNP1 | pSS1616 | F2 | R1 or S2 |
| pFA6a-mIAA7-ALFA-HIS3MX6 | pSS1633 | F2 | R1 or S2 |
| pFA6a-mIAA7-ALFA-kITRP1 | pSS1634 | F2 | R1 or S2 |
| pFA6a-mIAA7-ALFA-kanNP1 | pSS1635 | F2 | R1 or S2 |

**Table S4.** Integrative expression plasmids for degron receptors. For genomic integration, expression plasmids need to be linearised by restriction digest.

| Plasmid | Alias | Restriction digest with |
| --- | --- | --- |
| pRS405-P<sub>ADH</sub>-OsTIR1(F74G) | pSS1330 | AgeI or EcoRI |
| pRS406-P<sub>ADH</sub>-OsTIR1(F74G) | pSS1331 | StuI or BstBI |
| pNH605-P<sub>ADH</sub>-OsTIR1(F74G) | pSS1354 | PmeI |
| pNH604-P<sub>ADH</sub>-OsTIR1(F74G) | pSS1361 | PmeI |
| pNH605-P<sub>ADH</sub>-OsTIR1 | pSS1402 | PmeI |
| pNH605-P<sub>ADH</sub>-OsTIR1(F74A) | pSS1444 | PmeI |
| pRS405-P<sub>ADH</sub>-AtAFB2 | pSS1195 | AflII or AjuI |
| pNH605-P<sub>ADH</sub>-AtAFB2 | pSS1446 | PmeI |
| pNH605-P<sub>ADH</sub>-AtAFB2(F74G) | pSS1447 | PmeI |
| pNH605-P<sub>ADH</sub>-AtAFB2(F74A) | pSS1448 | PmeI |
| pRS303K-P<sub>GPD</sub>-OsTIR1(F74G) | pSS1553 | NheI/AfeI (double digest) |
| pRS303K-P<sub>ADH</sub>-OsTIR1(F74G) | pSS1554 | NheI/AfeI (double digest) |
| pRS303K-P<sub>RNR1</sub>-OsTIR1(F74G) | pSS1555 | NheI/AfeI (double digest) |
| pRS303H-P<sub>ADH</sub>-OsTIR1(F74G) | pSS1631 | NheI/AfeI (double digest) |
| pRS303N-P<sub>ADH</sub>-OsTIR1(F74G) | pSS1632 | NheI/AfeI (double digest) |
| pRS303K-P<sub>GAL</sub>-OsTIR1(F74G)-GEM | pSS1624 | NheI/AfeI (double digest) |

